# Single-cell and spatial transcriptomics reveal the pathogenesis of chronic granulomatous disease in a natural model

**DOI:** 10.1101/2025.01.23.634507

**Authors:** Hanzhi Yu, Guorong Zhang, Yunxi Ma, Jiayu Ding, Jingjing Liu, Zhiliang Zhou, Shaozhuo Jiao, Ge Dong, Zhigang Cai

## Abstract

Genetic defects in NOX2 can cause chronic granulomatous disease (CGD), characterized by increased susceptibility to infections and pronounced inflammatory responses that lead to granuloma formation. We developed a CGD model using *Ncf2*^-/-^ mice through controlled environmental exposure. Unlike in specific pathogen-free environment, these mice spontaneously developed pulmonary granulomas under clean-grade conditions. Employing single-cell RNA sequencing, we observed an expansion of neutrophils and monocyte-derived macrophages (MDMs) within the lung tissue, identifying pro-inflammatory NOS2^high^-neutrophils and a distinct MDMs subset marked by NOS2 and ARG1. Spatial transcriptomics demonstrated that NOS2^high^-neutrophils were located at the core area, while the MDMs subset was positioned at the periphery, facilitating extracellular matrix remodeling. Pharmacological targeting of MIF, deletion of the pro-survival gene *Morrbid*, and elimination of *Il1r1* all suppressed granuloma formation by mitigating inflammation. This study underscores how environmental control can establish a natural CGD model, elucidates the mechanisms of granuloma formation, and develops potent therapeutic interventions.

**Highlights:**

1. Through installing a mutation in a NOX2 subunit gene *Ncf2* and changing the husbandry environment, we generated a natural CGD mouse model with progressive pulmonary granuloma formation.
2. NOS2^high^ neutrophils located in the core region of CGD lung granulomas, exhibit a pro-inflammatory transcriptional profile.
3. Monocyte-derived macrophages marked by *Nos2* and *Arg1* are located at the periphery of granulomas and exhibit a profibrotic transcriptional profile.
4. The transcriptomic studies assist the development of three effective perturbations for treating the rare disease CGD via targeting *Il1r1, Morrbid,* and *Mif* respectively.

## INTRODUCTION

Chronic granulomatous disease (CGD) is a primary immunodeficiency caused by loss-of-function mutations in the genes encoding the NADPH Oxidase 2 (NOX2) subunits (*CYBB, CYBA, NCF1, NCF2, NCF4*) and the subunit assembly-related gene (*CYBC1*).^1^ These genetic variants impair the production of reactive oxygen species (ROS) by innate immune cells such as neutrophils, monocytes, and macrophages, which are essential for killing microbes. Consequently, this impairment leads to severe and recurrent bacterial and fungal infections, with pulmonary infections being particularly prevalent and the primary cause of mortality.^2–4^ Besides infection, the loss of ROS production also halts the inflammatory response, leading to severe inflammatory complications such as granuloma formation, inflammatory bowel disease, and systemic lupus erythematosus.^4,5^ Effective controlling inflammation is crucial for reducing tissue damage and autoimmune phenomena, as well as for improving the success rates of transplantation and gene therapy for CGD.^6,7^

In CGD patients and some animal model mimicking CGD with specific pathogen challenging, the proliferation and mobilization of myeloid immune cells, especially neutrophils, coupled with the significant expression of pro-inflammatory cytokines such as TNF-α, IL-1α, and IL-1β, are considered major contributors to the induction of inflammatory states. Immunomodulatory strategies that target these mediators, including the TNF-α inhibitor Infliximab and the IL-1 receptor antagonist Anakinra, have been used to treat related inflammatory complications with some success.^8,9^

However, these treatments are constrained by substantial side effects or low remission rates in severely ill patients. Therefore, constructing a comprehensive immune cell atlas of CGD to understand the key endogenous signals driving persistent inflammation in the special disease is critical for developing new targeted therapies.

Macrophage migration inhibitory factor (MIF) is a significant regulator in inflammatory and autoimmune diseases, known to intensify inflammation by activating signaling pathways such as MAPK, PI3K/AKT, and NF-κB.^10^ However, the role of MIF in CGD remains unexplored and represents a potential area for further research. Furthermore, timely apoptosis and clearance of neutrophils are crucial for terminating the inflammatory response. Under the influence of inflammatory cytokines such as TNF-α, IL-1, and IL-8, apoptosis of neutrophils is delayed, leading to increased tissue damage and inflammation.^11^ *Morrbid* (Myeloid RNA Regulator of Bim Induced Death) is a long non-coding RNA (lncRNA) specifically expressed in myeloid lineage that can prolong cell survival by downregulating the expression of Bim, a protein associated with apoptosis.^12^ However, persistent expression of *Morrbid* may result in the excessive survival of myeloid cells, further exacerbating tissue damage. In our previous study, we have demonstrated that loss of *Morrbid* mitigate myeloid-lineage leukemogenesis. Here we hypothesize that *Morrbid* plays a role in driving excessive inflammatory state in CGD.

In our most recent study, we demonstrated that *NCF2* is associated with acute myeloid leukemia and anti-leukemia drug resistance.^13^ However, its physiological roles remain largely unknown. Neutrophil Cytosolic Factor 2 (*NCF2*) encodes the cytosolic subunit p67phox of the NOX2 holoenzyme, which is critical for the NADPH oxidase activity. Germline mutations in *NCF2*, as with other NOX2 subunit genes, have been reported to be associated with CGD.^14^ Notably, certain genetic variants of *NCF2* are also linked to increased susceptibility to autoinflammatory diseases such as systemic lupus erythematosus^15^ and rheumatoid arthritis^16^, underscoring its pivotal role in regulating intrinsic inflammation. In this study, we changed the husbandry condition from specific pathogen free facility (SPF) to a clean-grade (CL) condition and observed a natural occurrence of CGD in *Ncf2^-/-^* mice. We thoroughly decoded the pathogenies of the nature CGD by combining regular single-cell transcriptomics and spatial transcriptomics and developed three different but correlated perturbations for the rare hypo-inflammation condition.

## RESULTS

### *Ncf2^-/-^* mice spontaneously develop pulmonary granulomas under the CL condition

Hematoxylin and Eosin (H&E) staining and Masson’s trichrome (Masson) staining were employed to observe the pathological changes in the lung tissues of WT and *Ncf2^-/-^* mice raised in SPF and CL environments. H&E staining revealed that the lung tissues of SPF WT and CL WT mice were structurally intact: neatly arranged cells, clear alveolar walls, and no obvious infiltration of inflammatory cells (**Figure 1A**). SPF *Ncf2^−/−^* mice appear grossly normal, however, the lung tissue of CL *Ncf2^−/−^* mice exhibited almost complete consolidation, characterized by extensive infiltration of inflammatory cells and significant tissue necrosis (**Figure 1A, Figure S1A**). Masson staining further revealed that the infiltrated inflammatory cells were encased by blue collagen fibers, forming granulomatous structures (**Figure 1B**). In contrast, SPF *Ncf2^-/-^*mice exhibited no indications of infection or pulmonary granuloma formation (**Figure 1A**), suggesting that the pathogens present in the CL environment are very likely responsible for the lung infections and spontaneous granuloma manifestations.

**Figure 1.**
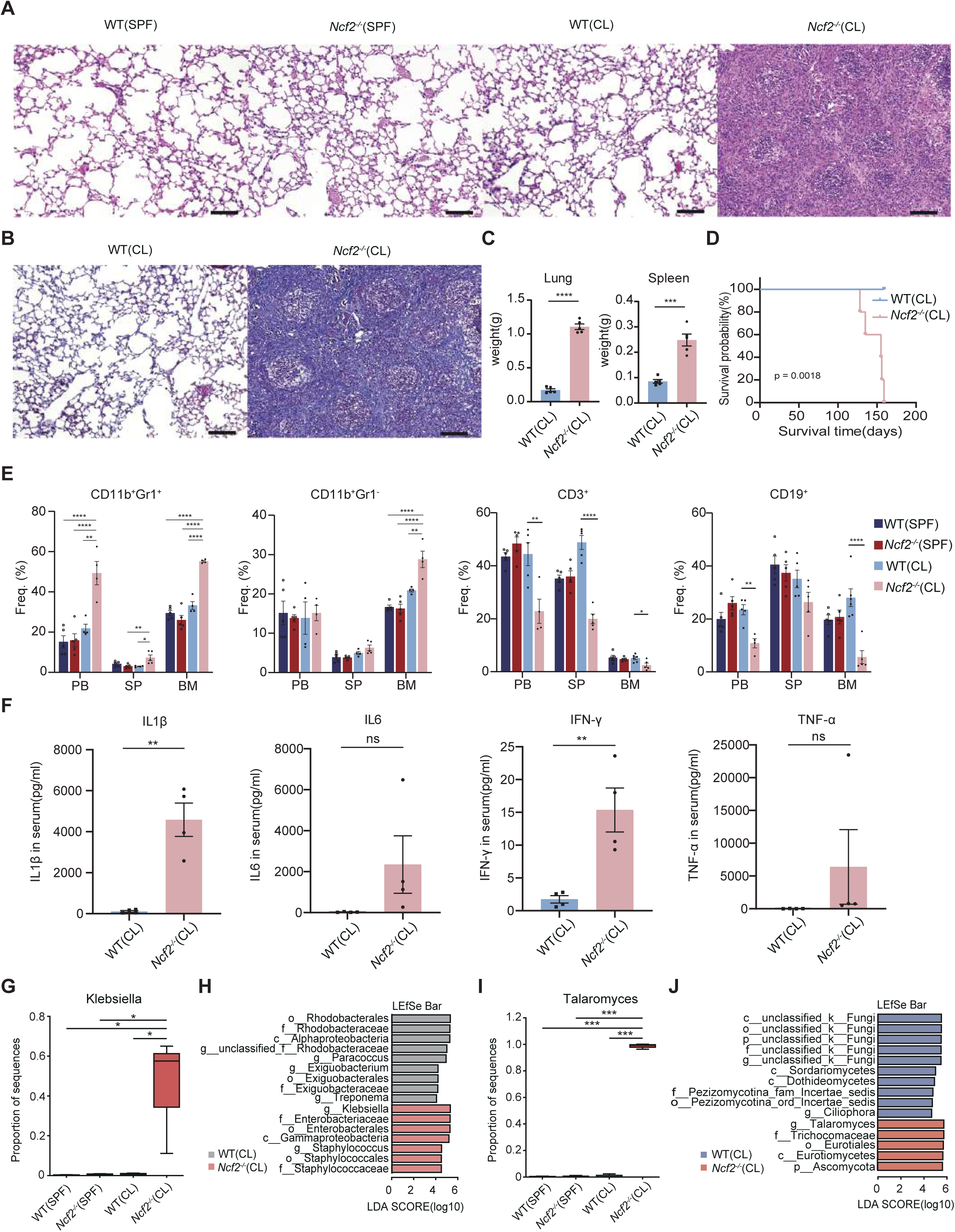
Constructing a CGD infection model through exposure to clean-grade environment **(A)** Hematoxylin and Eosin (H&E) staining of lung tissues from SPF WT, SPF *Ncf2*^-/-^, CL WT, and CL *Ncf2*^-/-^ mice. Scale bar: 100μm. **(B)** Masson’s trichrome staining of lung tissues from CL WT and CL *Ncf2*^-/-^ mice. Scale bar: 100μm. **(C)** Weight of lung and spleen organs from CL WT and CL *Ncf2*^-/-^ mice. **(D)** Survival curves of CL WT and CL *Ncf2*^-/-^ mice. **(E)** Proportions of myeloid and lymphoid cells in the peripheral blood (PB), spleen (SP), and bone marrow (BM) from SPF WT, SPF *Ncf2*^-/-^, CL WT, and CL *Ncf2*^-/-^ mice, revealed by flow cytometry. CD11b^+^Gr1^+^ for granulocytic populations; CD11b^+^Gr1^-^ for monocytic populations. **(F)** Protein levels of IL1β, IL6, IFN-γ, TNF-α in the serum of CL WT and CL *Ncf2^-/-^* mice determined by ELISA. **(G)** The abundance differences of the Klebsiella across the four groups. **(H)** LEfSe (Linear discriminant analysis Effect Size) analysis reveals differences in the pulmonary bacterial communities between CL WT and CL *Ncf2*^-/-^ mice. **(I)** The abundance differences of the Talaromyces across the four groups. **(J)** LEfSe analysis revealing differences in the pulmonary fungal communities between CL WT and CL *Ncf2*^-/-^ mices.* p < 0.05, ** p < 0.01,*** p < 0.001,****p < 0.0001.

Patients with CGD suffer from impaired pathogen clearance due to diminished production of reactive oxygen species (ROS) in phagocytic cells like neutrophils and macrophages, resulting in persistent and recurrent infections. The fluorescent probe DCFH-DA was used to evaluate the ROS levels in myeloid cells from the peripheral blood of CL WT and CL *Ncf2^-/-^* mice. Compared with CL WT mice, ROS production in myeloid cells of CL *Ncf2^-/-^*mice was significantly reduced, which is consistent with the immunodeficiency observed in *Ncf2^-/-^* mice (**Figure S2A**).

The lung and spleen weights of CL *Ncf2*^-/-^ mice were markedly greater than those of CL WT mice, attributable to persistent lung infections and augmented extramedullary hematopoiesis (**Figure 1C**). This also compromised the survival of the mice, resulting in a significant reduction in the lifespan of CL *Ncf2*^-/-^ mice (**Figure 1D**, survival range: 128∼159 days). Given the marked infection manifestations in CL *Ncf2^-/-^*mice, we utilized flow cytometry to further investigate alterations in the immune cell composition within the peripheral blood (PB), spleen (SP), and bone marrow (BM). Compared to SPF WT mice, SPF *Ncf2^-/-^* mice exhibited no significant alterations in the proportions of myeloid cells (CD11b^+^GR1^+^granulocytic populations and CD11b^+^GR1^-^monocytic populations) and lymphoid cells (CD3^+^ T lymphocytes and CD19^+^ B lymphocytes) in the PB, SP, and BM (**Figure 1E, Figure S3A and 3B**). These results further indicate that SPF barrier facilities protect *Ncf2*^-/-^ mice from infection and the associated mobilization of immune cells. However, compared to CL WT mice, CL *Ncf2^-/-^* mice exhibited an expansion of myeloid components and manifested a significant increase in CD11b^+^GR1^+^ granulocytes in the PB, SP, and BM, and a significant increase in CD11b^+^GR1^-^ monocytic populations in the BM (**Figure 1E**, **Figure S3A**). Accordingly, there was a reduction in lymphoid components, characterized by a significant decrease in the proportion of CD3^+^ T lymphocytes in the PB, SP, and BM, and a significant decrease in CD19^+^ B lymphocytes in the PB and BM (**Figure 1F**, **Figure S3B**). Enzyme linked immunosorbent assay (ELISA) was used to measure the levels of inflammatory cytokines in serum. Compared to CL WT mice, CL *NCF2*^-/-^ mice exhibited dramatically elevated levels of IL-1β (fold change: 40.4; p < 0.01), IL-6 (fold change: 85.4; p = 0.1491), IFN-γ (fold change:8.9; p < 0.01), and TNF-α (fold change:251.0; p = 0.3069) (**Figure 1F**). These results demonstrated that the inflammatory cells were infiltrated in the lung, accompanied by systematic inflammatory storm and altered hematopoiesis in the mutant animal.

### Altered microbiota in the lung tissues of the *Ncf2^-/-^* mice

Given the significant infection in the lungs of CL *Ncf2^-/-^*mice, we explored the microbiota composition within GGD pulmonary granulomas. Employing 16S rRNA sequencing, we investigated the influence of the CL environment on the bacterial communities in the lung tissue of CGD mice. The Chao index and the Shannon index were utilized to assess the abundance and diversity of the microbial communities, respectively. The findings revealed a reduction in both bacterial abundance and diversity in *Ncf2^-/-^* samples compared to their wild-type counterparts under both SPF and CL conditions (**Figure S4A**). Subsequently, we assessed the bacterial microbiota composition in mouse lung tissue samples at the genus level (**Figure S4B**). The results indicated that Acinetobacter and Pseudomonas, both prevalent in the lower respiratory tract of healthy individuals, predominated in the SPF *Ncf2^-/-^*and SPF WT groups (**Figure S4B**). In CL *Ncf2^-/-^* mice, the most dominant bacterial genera were Klebsiella, Staphylococcus, and Pelomonas (**Figure S4B**). Remarkably, Klebsiella and Staphylococcus were nearly exclusive to the CL *Ncf2^-/-^* group (**Figure 1G** and **Figure S4C**). The LEfSe (Linear discriminant analysis Effect Size) analysis revealed a significant elevation in the abundance of Klebsiella and Staphylococcus in the lung tissues of CL *Ncf2^-/-^*mice compared to the CL WT mice (**Figure 1H**), further substantiating the heightened susceptibility of CGD mice to these prevalent CGD-related pulmonary pathogens. Employing ITS sequencing, our investigation delved into the effects of exposure to a CL environment on the fungal communities within the lung tissues of CGD mice. The results indicated a consistent decline in fungal abundance in *Ncf2^-/-^* mice across both SPF and CL conditions (**Figure S5A**). Additionally, fungal diversity in the CL *Ncf2^-/-^* samples was significantly reduced compared to CL WT samples (**Figure S5A**). At the genus level, the Ascomycota phylum occupied a significant proportion in both the WT and *Ncf2^-/-^* samples under SPF conditions (**Figure S5B**). However, in the CL *Ncf2*^-/-^mice, Talaromyces became the dominant fungal community (**Figure S5B**). Both Kruskal-Wallis tests and LEfSe analysis further confirmed a significant increase in the abundance of Talaromyces in the CL *Ncf2^-/-^* group (**Figure 1I** and **J**). Overall, *Ncf2^-/-^* mice spontaneously develop pulmonary granulomas under CL conditions due to environmental pathogens, independent of Aspergillus spores, Candida albicans, Staphylococcus aureus, zymosan or other stimuli as suggested by the previous reports.^17–20^ This observation offers a new perspective for constructing a natural CGD model.

### Neutrophils and Monocyte-derived macrophages (MDMs) significantly accumulated in CGD mice lung tissue during infection

We conducted single-cell RNA sequencing (scRNA-seq) analysis of lung tissues from SPF WT, SPF *Ncf2^-/-^*, CL WT, and CL *Ncf2^-/-^* mice to reveal the transcriptional alterations of immune and stromal cells at the single-cell level. Following rigorous quality control, a total of 25,296 high-fidelity sequenced cells were obtained for further analysis, with each cell containing an average of 6,466 mapped reads and 1,972 genes. Cells were subjected to unsupervised clustering based on the differential expression of marker genes and visualized using Uniform Manifold Approximation and Projection (UMAP) (see Methods for details). Referring to lineage-specific genes reported in prior studies,^21–23^ we manually annotated cell clusters within our scRNA-seq datasets (**Figure 2A**, **Figure S6A**). We identified nine immune cell types in the lung tissue, including monocytes, macrophages, dendritic cells, neutrophils, T cells, natural killer cells, innate lymphoid cells, B cells, and plasma cells. Additionally, a cluster characterized by high expression of cell cycle-related genes (*Mki67* and *Top2a*) was identified as a mix of various immune cell types. We also identified nine stromal cell types, including type I and type II alveolar epithelial cells, ciliated cells, goblet cells, endothelial cells, fibroblasts, pericytes, mesothelial cells, and glial cells.

**Figure 2.**
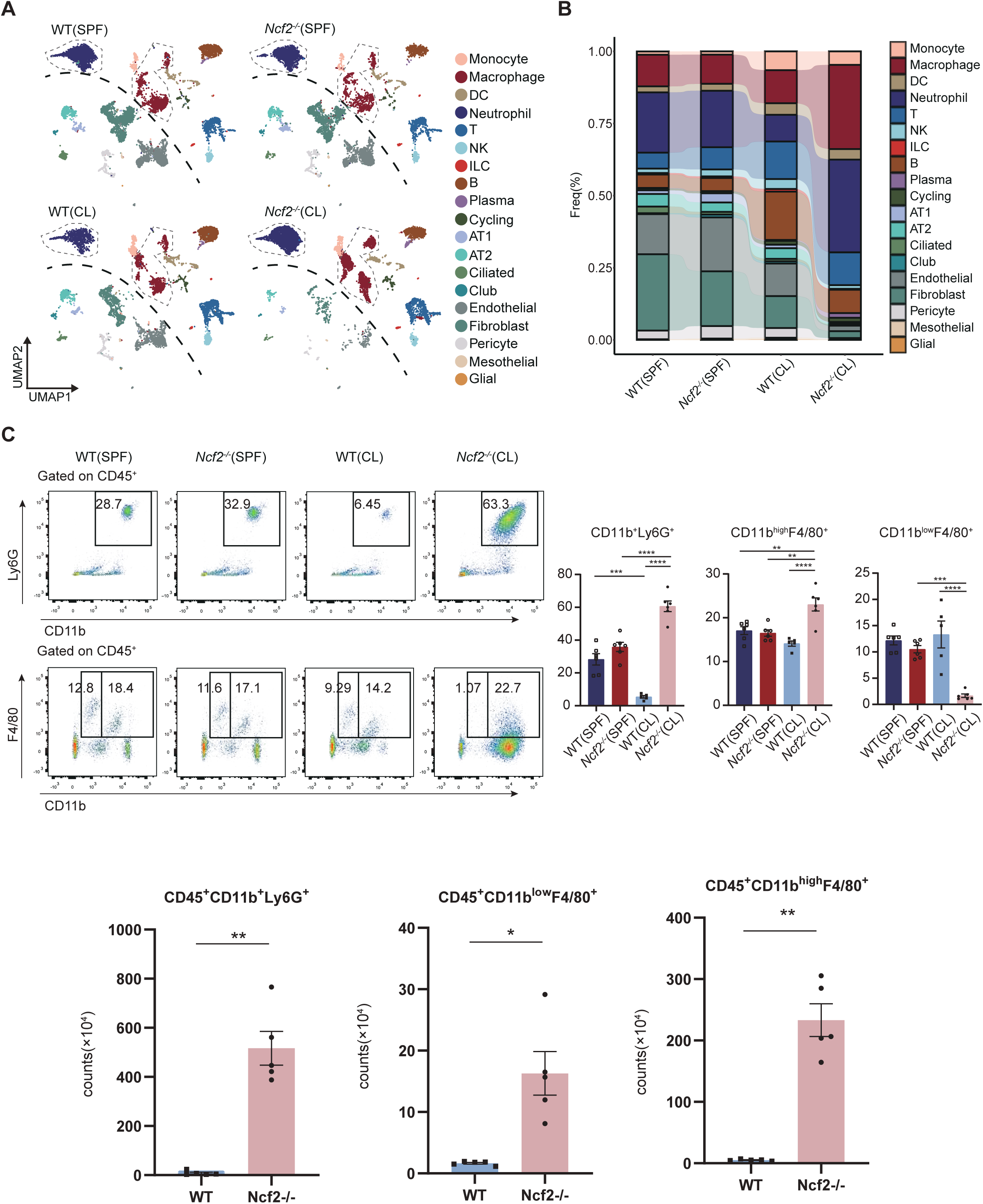
Analysis of immune landscape in CGD mice lung at single-cell resolution **(A)** The lung tissues were subjected for the single-cell RNA sequencing (scRNA-seq) analysis and UMAP plots reveals the distinct immune and stromal cell types in the lungs from SPF WT (n=11,305), SPF *Ncf2*^-/-^ (n=7,948), CL WT (n=7,912), and CL *Ncf2*^-/-^ (n=8,131) mice. In total 18 kinds of cell-type were annotated and colored across four scenarios. DC: dendritic cells, ILC: innate lymphoid cells, AT1: type I alveolar epithelial cells, AT2: type II alveolar epithelial cells, UMAP: uniform manifold approximation and projection. **(B)** Stacking bar plots showing the proportion of each cell-type cross four conditions. **(C)** Representative flow cytometry profiles of neutrophils and macrophages from lung tissues of SPF WT, SPF *Ncf2*^-/-^, CL WT, and CL *Ncf2*^-/-^ mice (left-up and left-down panel). Bar plots display the quantification of the proportions of CD11b^+^Ly6G^+^ neutrophils, CD11b^high^F4/80^+^ monocyte-derived macrophages (MDMs), and CD11b^low^F4/80^+^ resident macrophages (right panel). ** p < 0.01, *** p < 0.001, ****p < 0.0001.

The UMAP plots shows that immune cell clusters in the upper right of the black dashed line, while stromal cells gather in the lower left, highlighting the transcriptional heterogeneity between immune and stromal cells. Compared to SPF WT mice, SPF *Ncf2^-/-^* mice exhibited no notable changes in the fractions of immune and stromal cells (**Figure 2B**), consistent with the above observations that SPF *Ncf2^-/-^* mice appear grossly normal (**Figure 1A** and **Figure** S**1A**). However, CL *Ncf2^-/-^* mice revealed an increase in fractions of myeloid cells (neutrophils, macrophages) and a decrease in fractions of lymphocytes (T cells, B cells, NK cells) compared to that in CL WT mice (**Figure 2B**). Moreover, stromal cells, including type I alveolar epithelial cells, type II alveolar epithelial cells, endothelial cells, fibroblasts, and pericytes, were significantly diminished in CL *Ncf2^-/-^* mice compared to that in CL WT mice (**Figure 2B**), indicating a possible compromise of stromal cells during infection and pulmonary granuloma formation.

Among the immune cell subgroups, neutrophils and macrophages exhibited the most pronounced changes. We further employed flow cytometry analysis to assess these alterations observed from the scRNA-seq datasets. The flow cytometry analysis validated that environmental changes significantly affect the immune composition of lung tissue in WT mice, with a notably higher proportion of neutrophils in SPF WT mice compared to CL WT mice (**Figure 2C**). This phenomenon, also reported by Weiss et al^24^, may be attributed to enhanced lung basal immunity caused by highly sterile SPF barrier facilities. Under SPF conditions, the proportion of neutrophils in the lung tissues of *Ncf2^-/-^* mice showed no significant changes compared to WT mice. However, under CL conditions, there was a dramatic increase in the proportion of neutrophils in the lung tissues of *Ncf2^-/-^* mice (**Figure 2C**). Previous studies have established that CD11b^high^F4/80^+^ pulmonary macrophages are MDMs, while CD11b^low^F4/80^+^ pulmonary macrophages are predominantly resident macrophages (including alveolar macrophages and interstitial macrophages). Employing this flow cytometry strategy^25^, we observed a significant increase in CD11b^high^F4/80^+^ MDMs and a significant reduction in CD11b^low^F4/80^+^ resident macrophages in CL *Ncf2^-/-^* mice (**Figure 2C**, **Figure S7A**). This suggests that during the inflammatory response in CGD scenarios, resident macrophages may be depleted due to pathogen clearance, while this depletion can be compensated by the recruitment of MDMs from the BM.

### NOS2^high^ neutrophils were significantly expanded in the lung tissue of CGD mice

To investigate the role of neutrophils in CGD pulmonary granulomas, we performed dimensionality reduction and unsupervised clustering on the single-cell transcriptomic profiles of all neutrophils, yielding six distinct subgroups (namely Neu1 to Neu6) (**Figure 3A**). Notably, the Neu6 neutrophils were markedly elevated in the CL *Ncf2-/-* when compared to the other three scenarios, highlighting their critical role in the pulmonary immune response associated with CGD (**Figure 3B**). Differential gene expression analysis indicated that the transcriptional profiles of the Neu6 subset was characterized by the upregulation of pro-inflammatory genes such as *Nos2, Saa3*, and *Mif* (**Figure 3C**). Furthermore, Gene Ontology (GO) enrichment analysis revealed that inflammation-related signaling pathways, such as leukocyte chemotaxis, neutrophil activation, production of IL-1 and IL-6, and the NF-κB signaling pathway, were significantly enriched in Neu6 (**Figure 3D**). These findings underscore the pivotal role of Neu6 in mediating the inflammatory response, particularly through the production of inflammatory mediators and chemotactic migration. Simultaneously, excessive accumulation of Neu6 may also contribute significantly to tissue damage during the granuloma formation.

**Figure 3.**
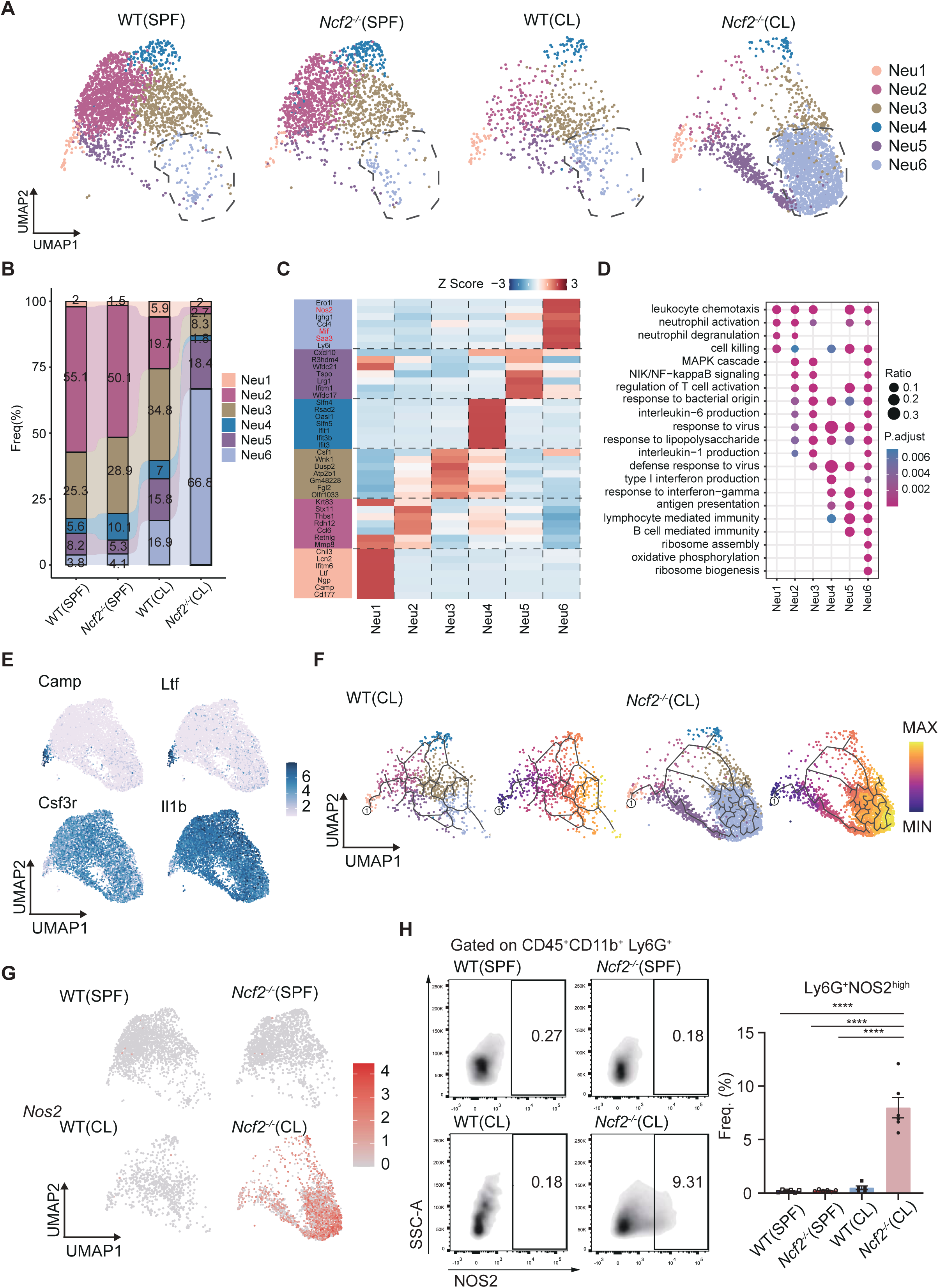
Transcriptional alterations in the neutrophil compartment of CGD mice **(A)** UMAP plots displaying sub-clusters of neutrophils in the lungs of SPF WT, SPF *Ncf2*^-/-^, CL WT, and CL *Ncf2*^-/-^ mice. **(B)** Stacking bar plots illustrating the frequency of distinct neutrophil subsets across the four conditions. **(C)** Heatmap showing the top marker genes (Top7) for each lung neutrophil subsets. **(D)** Bubble plots for Gene Ontology (GO) analysis across all neutrophil subsets. **(E)** Expression of indicated genes for labeling immature (*Camp*, *Ltf*) and mature (*Csf3r*, *Il1b*) neutrophils. **(F)** Pseudo-time trajectory of the neutrophil compartments analyzed by Monocle3. **(G)** Expression of *Nos2* in pulmonary neutrophils. **(H)** Flow cytometry profiles of NOS2^high^ neutrophils (left panel) in the lung tissue of SPF WT, SPF *Ncf2*^-/-^, CL WT, and CL *Ncf2*^-/-^ mice, with statistical analysis with bar plots (right panel).

As shown in **Figure 3E**, expression of marker genes in the UMAP plots illustrates that Neu1 sub-cluster distinctly expressed granulocyte genes characteristic of primitive or immature neutrophils, such as *Camp* and *Ltf*. In contrast, sub-clusters Neu2-6 exhibited expression of genes associated with mature neutrophils, including *Csf3r* and *Il1b*. Subsequently, the cell trajectory algorithm Monocle3 was employed to perform pseudo-time analysis across CL WT and CL *Ncf2^-/-^* sample, aiming to delineate the differentiation trajectories within the neutrophil compartment. Starting with the immature sub-cluster Neu1, the analysis revealed that the Neu6 subgroup, which increased in the CL *Ncf2^-/-^* mice, represents the most mature state of neutrophil differentiation (**Figure 3F**).

Furthermore, as shown in **Figure 3G**, the expression of marker genes in the UMAP plots indicates that the nitric oxide synthase 2 gene (*NOS2*), crucial for the production of nitric oxide (NO), is predominantly expressed in neutrophils from the CL *Ncf2^-/-^*group, with a significant enrichment in the Neu6 subset. Given that *Nos2* is also a top marker gene for the Neu6 cluster (**Figure 3C**), it was selected as a representative marker for this subset and validated through flow cytometry (**Figure 3H**). The flow cytometry results demonstrated a significant increase in NOS2^high^ neutrophils in the lung tissue of CL *Ncf2*^-/-^ mice, whereas only a small number of such cells were present in the other three groups (**Figure 3H**). This result suggests a distinctive role of NOS2^high^ neutrophils in the pulmonary inflammatory response within this specific genetic and environmental context, highlighting their potential significance in the pathology of CGD or general granuloma formation.

### MDMs in CGD mice exhibit both M1 and M2 polarization

In addition to neutrophils, macrophages underwent significant changes during granuloma formation in CGD mice. To further investigate the transcriptional changes and differentiation processes in such cells, we re-clustered the monocyte and macrophage compartments, identifying five distinct sub-clusters: Mono1, Mono2, Mac1, Mac2, and Mac3 (**Figure 4A** and **B**). Mono1 is distinguished by high expression of *Ly6c2*, representing the classical monocyte subgroup (**Figure 4C**). Mono2, characterized by low *Ly6c2* and high *Ace* expression, defined non-classical monocytes (**Figure 4C**). The Mac1 subset prominently expressed the M1 marker *Nos2*, the M2 marker *Arg1*, the inflammatory factor serum amyloid A3 (*Saa3*), as well as the extracellular matrix (ECM) remodeling genes matrix metalloproteinase 12 (*Mmp12*) and secreted phosphoprotein 1 (*Spp1*) (**Figure 4C**). Mac2 features genes linked to interstitial lung macrophages, such as *Cx3cr1*, *Folr2*, and *Lyve1* (**Figure S8A**). Additionally, highly mRNA expression of *Ear1, Plet1*, and *Siglecf* in the Mac3 subgroup identified it as an alveolar macrophage subgroup (**Figure 4C** and **Figure S8A**). Notably, the Mac1 subset was markedly increased in the lung tissue of the CL *Ncf2*^-/-^ mice and shows continuity relationship with classical monocytes (Mono1) on the UMAP plot (**Figure 4A** and **B**), suggesting that the Mac1 subgroup may arise from the differentiation of infiltrating monocytes.

**Figure 4.**
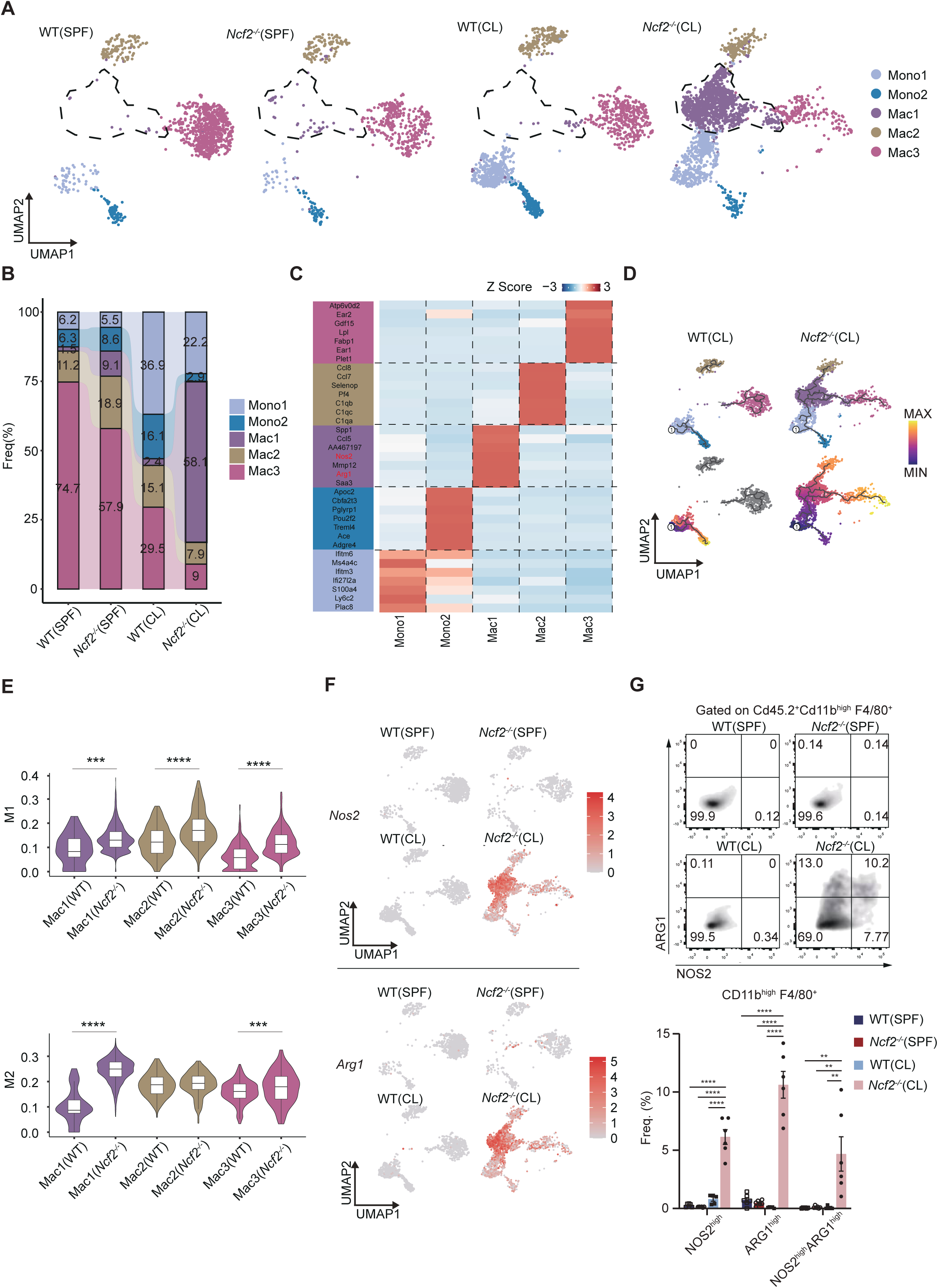
Transcriptional alterations in the monocyte-macrophage compartment of CGD mice **(A)** UMAP plots displaying sub-clusters of the monocyte-macrophage compartment in the lungs of SPF WT, SPF *Ncf2*^-/-^, CL WT, and CL *Ncf2*^-/-^ mice. **(B)** Stacking bar plots illustrating the frequency of distinct monocyte/macrophage subsets across the four conditions. **(C)** Heatmap showing the top marker genes (Top7) for each monocyte/macrophage subsets. **(D)** Pseudotime trajectory of the monocyte-macrophage compartment analyzed using Monocle3. **(E)** Violin plot displaying the M1/M2 module scores for macrophage subsets. **(F)** UMAP plots displaying the expression of M1 marker gene (*Nos2*) and M2 marker gene (*Arg1*) in pulmonary MDMs. **(G)** Flow cytometry analysis of NOS2^high^, ARG1^high^, and NOS2^high^ARG1^high^ MDMs in the lung tissue of SPF WT, SPF *Ncf2*^-/-^, CL WT, and CL *Ncf2*^-/-^ mice (left panel), with statistical data depicted in the bar plots (right panel).

To further explore the connections between monocyte and macrophage clusters, we employed the Monocle3 algorithm to infer the dynamic immune states and cell transition processes within the monocyte-macrophage lineage, as illustrated in **Figure 4D**. Given the classical monocytes’ capability to migrate into tissues and differentiate into various macrophage types, we designated them as the starting point of the trajectory. In the analyses of CL *Ncf2*^-/-^ mice, we observed that the resident macrophages Mac2 and Mac3 were situated at the end of the trajectory, whereas the Mac1 cluster was located at the center position, mediating the differentiation of classical monocytes into resident macrophages (**Figure 4D**). Accordingly, we considered Mac1 as monocyte-derived macrophages. In contrast, the absence of the Mac1 subgroup in CL WT mice led to truncated pseudo-time trajectories, suggesting that the interstitial macrophage Mac2 and alveolar macrophage Mac3 subgroups may primarily rely on self-renewal for replenishment (**Figure 4D**). These results highlighted the significant contribution of monocytes to the differentiation of resident macrophages under inflammatory conditions, emphasizing the crucial role of the Mac1 subgroup in bridging this transformation process.

For a long time, macrophages have been categorized into pro-inflammatory M1 and anti-inflammatory M2 types based on *in vitro* stimulation experiments.^26^ M1 macrophages contributed to tissue damage through the secretion of pro-inflammatory cytokines, while M2 macrophages mitigated inflammatory responses and promoted tissue repair by expressing anti-inflammatory factors. We utilized a list of M1 and M2 genes to assess the M1 or M2 polarization scores within the macrophage compartment. Violin plots demonstrated a significant elevation in M1 polarization scores across all lung macrophage subsets in CL *Ncf2*^-/-^ mice, accompanied by increased M2 polarization scores in Mac1 and Mac3 (**Figure 4E**). UMAP visualization revealed that the M1 marker *Nos2* and the M2 marker *Arg1* were exclusively expressed in CL *Ncf2*^-/-^ macrophages and were particularly enriched in the Mac1 subgroup (**Figure 4F**). Furthermore, considering *Nos2* and *Arg1* as top marker genes for the MAC1 subgroup (**Figure 4C**), we chose *Nos2* and *Arg1* as representative markers for this subgroup. Flow cytometry analysis revealed a significant expansion of specific NOS2^high^, ARG1^high^, and NOS2^high^ARG1^high^ MDMs in the pulmonary tissue of CL *Ncf2*^-/-^ mice, while these cells were virtually absent in other three scenarios (**Figure 4G**).

### Spatial transcriptomic analysis reveals the structural and transcriptional characteristics of CGD pulmonary granulomas

In order to analyze the molecular and cellular architecture of CGD pulmonary granuloma, we employed a single-nuclei level spatial transcriptome platform (SeekSpace, SeeGene BioScience, See Methods for details) to analyze the lung tissue slices from CL WT and CL *Ncf2*^-/-^ mice. This technique is capable of capturing independent nuclei, achieving spatial precision at the single-cell (nuclei) level. The robust cell type decomposition (RCTD)^27^ algorithmwas utilized to precisely annotate cell positions within the spatial transcriptomics data based on annotation information from the single-cell dataset (see Methods for detail). In the CL WT section, goblet cells were predominantly located in the bronchial region, while alveolar epithelial cells, monocytes, and macrophages were dispersed throughout the lung tissue without regional specificity (**Figure 5A** and **B**). In the CL *Ncf2*^-/-^ sample, a large granulomatous necrotic core was observed, where capturing a single cell was nearly impossible due to nuclear dissolution and fragmentation (**Figure 5A** and **B**). Around the necrotic core, there was an enrichment of monocytes and macrophages, along with a few neutrophils (**Figure 5B**). Macrophages, being the predominant immune-regulatory cell type forming the granuloma, were considered key drivers of granuloma formation.^28^ Notably, in the spatial transcriptomics data of CL *Ncf2*^-/-^, the specific macrophage subgroup Mac1, identified in the single-cell data, was accurately annotated and predominantly located at the margins of the granulomas (**Figure 5C**). We further analyzed the expression of the marker genes *Arg1* and *Nos2* across all macrophages and specifically within the Mac1. The findings revealed that the expression of *Arg1* and *Nos2* was predominantly concentrated in Mac1 (**Figure 5D**), further suggesting that these genes serve as indicative markers for this subgroup. Notably, The Mac1 subgroup exhibited elevated expression of *Fn1* (**Figure 5E**), which was crucial for constructing the extracellular matrix (ECM) framework. Moreover, the expression of *Mmp12* was also elevated in the Mac1 (**Figure 5E**). This enzyme was responsible for the degradation of extracellular matrix (ECM) components such as collagen, laminin, and fibronectin, indicating a potential role in promoting ECM remodeling and tissue fibrosis.^29^ Beyond immune cells, there was a notable enrichment of fibroblasts and vascular endothelial cells at the periphery of granulomas (**Figure 5B**). During granuloma formation, fibroblasts accelerated the fibrotic process in response to inflammatory stimuli, while vascular endothelial cells were essential for providing blood and lymphatic support.^30^ In the CL *Ncf2*^-/-^ mice, fibroblasts were demonstrated to highly express ECM component genes (*Fn1, Col1a1*) surrounding the necrotic core of the granulomas (**Figure 5F**).

**Figure 5.**
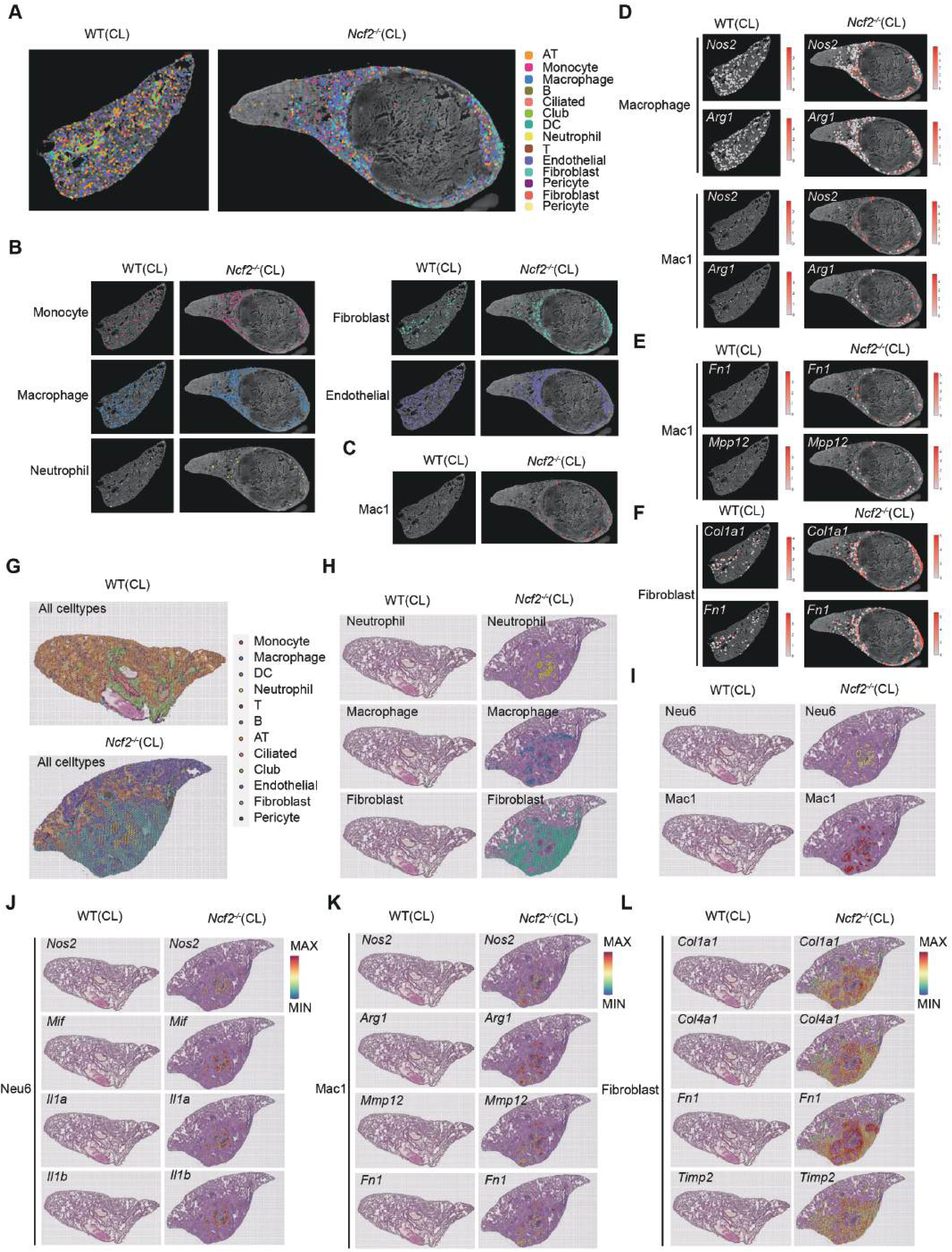
Decoding CGD granuloma structures using two different spatial transcriptomic approaches **A-F**, the results from the SeekSpace platform; G-L, the results from the S1000 platform **(A)** Spatial localization of main immune and stromal cell types in lung tissue sections of CL WT and CL *Ncf2*^-/-^ mice, identified via regular scRNA-seq datasets-assisted deconvolution RCTD. **(B)** Spatial localization of monocytes, macrophages, neutrophils, fibroblasts, and endothelial cells in lung sections of CL WT and CL *Ncf2*^-/-^ mice. **(C)** Spatial localization of the Mac1 subset in lung sections of CL WT and CL *Ncf2*^-/-^mice. **(D)** Spatial distribution and expression of selected marker gene (*Nos2*、*Arg1*) in all macrophages and the Mac1 subset. **(E)** Spatial distribution and expression of selected extracellular matrix (ECM) remodeling gene (*Fn1 Mmp12*) in the Mac1 subset. **(F)** Spatial distribution and expression of of selected ECM remodeling gene(*Fn1 Col1a1*) in fibroblasts. **(G)** Spatial localization of main immune and stromal cell types in lung tissue sections of CL WT and CL *Ncf2*^-/-^ mice, identified via RCTD. **(H)** Spatial localization of neutrophils, macrophages, fibroblasts in lung tissue sections of CL WT and CL *Ncf2*^-/-^ mice. **(I)** Spatial localization of the Neu6 and the Mac1 subset in lung sections of CL WT and CL *Ncf2*^-/-^ mice. **(J)** Spatial distribution and expression of selected marker gene (*Nos2*) and proinflammatory gene *(Mif, Il1a, Il1b*) in the Neu6 subset. **(K)** Spatial distribution and expression of selected marker gene (*Nos2*, *Arg1*) and ECM remodeling gene (*Mmp12, Fn1*) in the Mac1 subset. **(L)** Spatial distribution and expression of of selected ECM remodeling gene (*Col1a1*, *Col4a1*, *Fn1*, *Timp2*) in fibroblasts.

Due to the failure in capturing cells located in the core granuloma region with single-nuclei spatial transcriptomics, we further employed the BMKMANU S1000 Spatial Transcriptome to extend our findings. Utilizing the same RCTD algorithm, spatial locations within sections were deconvoluted and annotated as the predominant cell types. In lung tissue sections from CL WT mice, goblet and ciliated cells were predominantly localized in the bronchial region, whereas alveolar cells were evenly distributed throughout the lung field (**Figure 5G**). In CL *Ncf2*^-/-^ mice, we observed the formation of granuloma lesions. This time we observed that necrosis was evident in the core of the lesions, predominantly populated by infiltrating neutrophils (**Figure 5H**). Further analysis suggested that neutrophils in the core region were dominated by Neu6 subsets identified in our scRNA-seq datasets (**Figure 5I**, and **Figure 3A**). Surrounding the necrotic core, at the periphery of the granulomas, macrophages and fibroblasts were detected, with macrophages specifically classified as the Mac1 subgroup **(Figure 5H)**. In the spatial transcriptomics datasets, the Neu6 subgroup in the lung tissues of *Ncf2*^-/-^ mice also exhibit high expression of the marker gene *Nos2*, along with pro-inflammatory genes *Mif, Il1a,* and *Il1b* (**Figure 5J**). Mac1 expressed elevated levels of the marker genes *Nos2* and *Arg1*. Additionally, these macrophages ware involved in the degradation and remodeling of the ECM through the expression of *Mpp9*, *Mpp12, Mpp14*, and *Fn1* (**Figure 5K**). Moreover, fibroblasts situated at the periphery of the lesion are excessively activated, expressing elevated levels of ECM component genes (*Col1a1, Col4a1, Fn1*) and the ECM regulatory factor *Timp2*, thereby facilitating the occurrence and remodeling of the pathological tissue granuloma (**Figure 5L**).

In summary, combining the analysis of regular single-cell transcriptomics and spatial transcriptomics, we demonstrated that the Neu6 neutrophil subgroup, marked by high levels of pro-inflammatory cytokines in CGD mouse lung tissues, occupied the central region of granulomas, very likely playing a crucial role in pathogen eradication and intensifying inflammatory responses. Concurrently, Mac1 macrophages and fibroblasts, located at the periphery of the granulomas, engaged in ECM remodeling and the development of fibrotic encapsulation.

### IL1 receptor blockade reduced excessive inflammatory response in CGD mice

The pro-inflammatory cytokine interleukin-1 (IL-1) is crucial in numerous physiological processes and mediates the inflammatory and immune responses to many common diseases.^31^ IL-1 manifests in two agonistic forms, IL-1α and IL-1β, both of which interact with the sole signaling IL-1 type 1 receptor (IL-1R1) to initiate cellular signaling. In scRNA-seq datasets, we observed a significant upregulation of *Il1*a and *Il1b* in neutrophils and macrophages of CGD mice (**Figure S9A**). To investigate whether IL-1 signaling promoted pulmonary inflammation in CGD mice modeled through CL-grade environmental exposure, we constructed *Il1r1^-/-^Ncf2^-/-^* (double knockout, DKO) mice. Compared to the CL *Ncf2^-/-^* group, the CL *Ncf2^-/-^Il1r1^-/-^* lung tissue exhibited significantly reduced granulocyte infiltration, with no apparent granuloma lesions forming (**Figure S9B**). Additionally, in the CL *Ncf2^-/-^Il1r1^-/-^* group, there was a significant reduction in the proportion of Ly6G^+^ neutrophils, while the number of CD11b^low^F4/80^+^ resident macrophages was restored (**Figure S9C**). Subsequently, flow cytometry was used to identify immune cell fractions in the PB, SP, and BM. The results indicated that the genetic deficiency of *Il1r1* significantly reduced the proportion of CD11b^+^Gr1^+^ granulocytes in the PB, SP, and BM of CGD mice (**Figure S9D**). Simultaneously, the proportions of CD3^+^ T lymphocytes and CD19^+^ B lymphocytes were restored (**Figure S9D**). Therefore, our results demonstrate that IL-1 signaling plays a crucial role in mediating hyperinflammation in CGD mice modeled under CL conditions.

### 4IPP treatment regulated hyperinflammation in CGD mice via the MIF/NLRP3/IL1**β** axis

Macrophage Migration Inhibitory Factor (MIF) is a homotrimeric protein that acts as a pleiotropic proinflammatory cytokine, involved in a range of pathophysiological processes such as leukocyte recruitment, immune response, cell proliferation, and tumor formation.^10^ In both the scRNA-seq and spatial transcriptomics datasets, we observed that the Neu6 subset expanded in CL *Ncf2^-/-^* mice highly expressed *Mif* (**Figure 3C and 5J**). Additionally, *Mif* expression was significantly upregulated in both neutrophils and macrophages of CGD mice compared to controls (**Figure 6A**). We hypothesized that MIF might play a crucial role in mediating the excessive inflammatory response in CGD. 4-Iodo-6-phenylpyridine (4IPP) is a specific suicidal substrate for MIF, capable of covalently binding to MIF and irreversibly inhibiting its biological activity. We treated WT and *Ncf2^-/-^*mice with 4IPP (1 mg/kg) or an equivalent volume of 3% DMSO (the solvent for 4-IPP, the vehicle control) daily, starting from the onset of environmental exposure and continued for one month (**Figure 6B**). H&E staining demonstrated that, compared to the *Ncf2^-/-^* (DMSO) group, 4IPP treatment significantly reduced immune cell infiltration in the lung tissue of CL *Ncf2^-/-^*mice, with no obvious consolidation or granuloma formation (**Figure 6C**). Flow cytometry analysis further revealed a sharp decrease of Ly6G^+^ neutrophils in the lung tissue of the 4IPP treatment group (**Figure 6D**). Additionally, the infiltration of CD11b^high^F4/80^+^ MDMs was reduced, while the proportion of CD11b^low^F4/80^+^ resident macrophages was restored (**Figure 6D**). In addition, we analyzed the effect of 4IPP treatment on BM and peripheral immune cell fractions by flow cytometry. The results showed that 4IPP treatment significantly reduced the proportion of CD11b^+^Gr1^+^ granulocytes in PB, SP, and BM of CL *Ncf2^-/-^* mice compared to the DMSO control, suggesting a decrease in myeloid cell mobilization due to systemic inflammation (**Figure 6E** and **Figure S10A**). Concurrently, there was an increase in the proportion of CD3^+^ T lymphocytes and CD19^+^ B lymphocytes in PB, SP, and BM, indicative of a restoration of non-myeloid hematopoiesis (**Figure 6E**).

**Figure 6.**
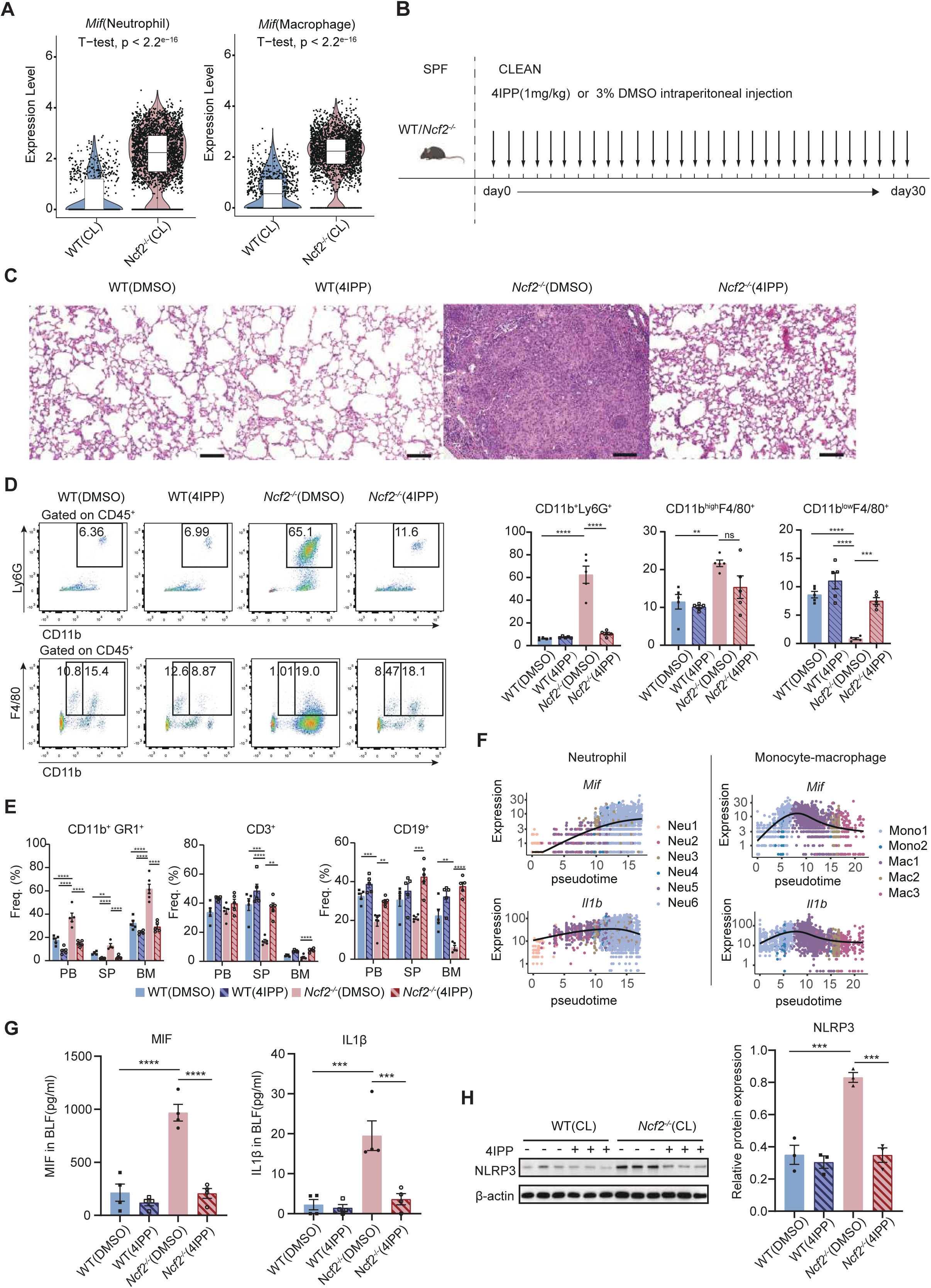
MIF inhibitor 4IPP alleviates hyperinflammation in CGD mice **(A)** Expression of *Mif* in neutrophils and macrophages in the scRNA-seq datasets. **(B)** Schematic diagram of the 4IPP treatment. **(C)** H&E staining of lung tissues from WT(DMSO), WT(4IPP), *Ncf2*^-/-^(DMSO), and *Ncf2*^-/-^(4IPP) mice. Scale bar: 100μm. **(D)** Representative flow cytometry plots of neutrophils and macrophages from lung tissues of WT(DMSO), WT(4IPP), *Ncf2*^-/-^(DMSO), and *Ncf2*^-/-^(4IPP) mice. Bar plots displaying the proportions of CD11b^+^Ly6G^+^ neutrophils, CD11b^high^F4/80^+^ MDMs, and CD11b^low^F4/80^+^ resident macrophages. **(E)** Proportions of granulocytes and lymphoid cells in the peripheral blood (PB), spleen (SP), and bone marrow (BM) of WT(DMSO), WT(4IPP), *Ncf2*^-/-^(DMSO), and *Ncf2*^-/-^(4IPP) mice, as analyzed by flow cytometry. **(F)** Scatter plot showing expression level of *Mif* and *Il1b* along with the pseudotime within the neutrophil and macrophage compartments. Colors represent different cell types. **(G)** MIF and IL1β protein levels in mice bronchoalveolar lavage (BAL) fluid determined by ELISA. **(H)** Expression of NLRP3 protein in mice lung tissue detected by Western blot.

Pseudo-timing analysis indicated that *Mif* and *Il1b* displayed analogous expression patterns in neutrophils and macrophages from CL *Ncf2^-/-^* mice, hinting at a possible regulatory interaction between these two factors (**Figure 6F**). Prior research has indicated that MIF is capable of regulating IL-1β release through the activation of the NLRP3 inflammasome.^32^ In the present study, ELISA was used to measure the protein levels of MIF and IL1β in bronchoalveolar lavage fluid (BALF) of WT(DMSO), WT(4IPP), *Ncf2^-/-^*(DMSO) and *Ncf2^-/-^*(4IPP) mice. As shown in **Figure 6G**, a significant increase in MIF levels in the BALF of *Ncf2^-/-^* (DMSO) mice were determined compared to that in WT (DMSO) mice, while 4IPP treatment significantly reduced MIF levels in the BALF of CL *Ncf2^-/-^* mice. Meanwhile, IL-1β exhibited a consistent expression pattern with MIF (**Figure 6G**). Western blot analysis revealed that, compared to that in the WT (DMSO) group, NLRP3 expression was significantly elevated in the lung tissue of *Ncf2^-/-^*(DMSO) mice, whereas 4IPP treatment significantly decreased NLRP3 expression (**Figure 6H and Figure S10B**). These findings suggest that the MIF/NLRP3/IL-1β signaling axis plays a crucial role in modulating the inflammatory response in CGD lungs.

### Systemic inflammation induced pronounced myeloid-biased hematopoiesis in CL *Ncf2*^-/-^ mice

Recently, numerous studies have suggested that chronic inflammation disrupts the normal balance of hematopoiesis, leading to myeloid-skewing, which expedites disease progression in a manner similar to a chain reaction.^33,34^ Since the significant expansion of myeloid cells and lymphopenia were observed in PB, SP and BM of CL *Ncf2*^-/-^ mice, as well as the elevated levels of inflammatory factors, we hypothesized that the myeloid-biased hematopoiesis takes place and play a role in CGD. Such aberrant hematopoiesis may further exacerbate inflammation by augmenting myeloid cell output. To validate such hypothesis, we employed scRNA-seq to analyze transcriptional alterations in hematopoietic precursor cells of both CL WT and CL *Ncf2^-/-^* mice.

Magnetic beads were employed to isolate linage-negative (Lin^-^) cells from the BM, enriching hematopoietic precursor cells for scRNA-seq. Following rigorous quality control, gene expression data from 19,215 cells (CL WT group: 9,687; CL *Ncf2*^-/-^ group: 9,528) were obtained for cluster analysis (**Figure 7A**). With reference to lineage-specific markers (**Figure S11A**), we manual annotated the hematopoietic and immune cells within the BM, identifying 15 distinct cell types. These included hematopoietic stem/progenitor cells (HSPC: *Kit, CD34, Egr1*), erythroid progenitor cells (ErP: *Cpox, Klf1*), megakaryocytic progenitor cells (Mkp; *Itga2b, Pf4*), granulocyte progenitor cells (Gp: *Gfi1, Elane*), common monocyte progenitor cells (cMop: *Csf1r, Ms4a3*) and dendritic cells (DC: *Itgax, Bst2, Siglech*), monocytes (*S100a4, Ccr2*), macrophages (*Adgre1, Fcgr1*), neutrophils (*S100a8, S100a9, Ly6g*), eosinophils (*Prg2, Epx, Ccr3*), basophils (*Fcer1a, Press34, Mcpt8*), natural killer cells (NK: *Gzma, Ncr1, Klre1*), T cells (T: *Cd3e, Lck, Trbc2*), B cells (B: *Cd79a, Cd19*), plasma cells (Plasma: *Mzb1, Jchain*) (**Figure 7A** and **S11A**). Stacked plots indicated an increased proportion of myeloid immune cells in the CL *Ncf2*^-/-^ group compared to the CL WT group, including neutrophils, macrophages, monocytes, and eosinophils (**Figure 7B**). This was consistent with flow cytometry results, where CD11b^+^ cells were significantly expanded in the BM of CL *Ncf2*^-/-^ mice, comprising over 80% of the total cells (**Figure 1E**), which may account for the reduced efficiency of bead sorting.

**Figure 7.**
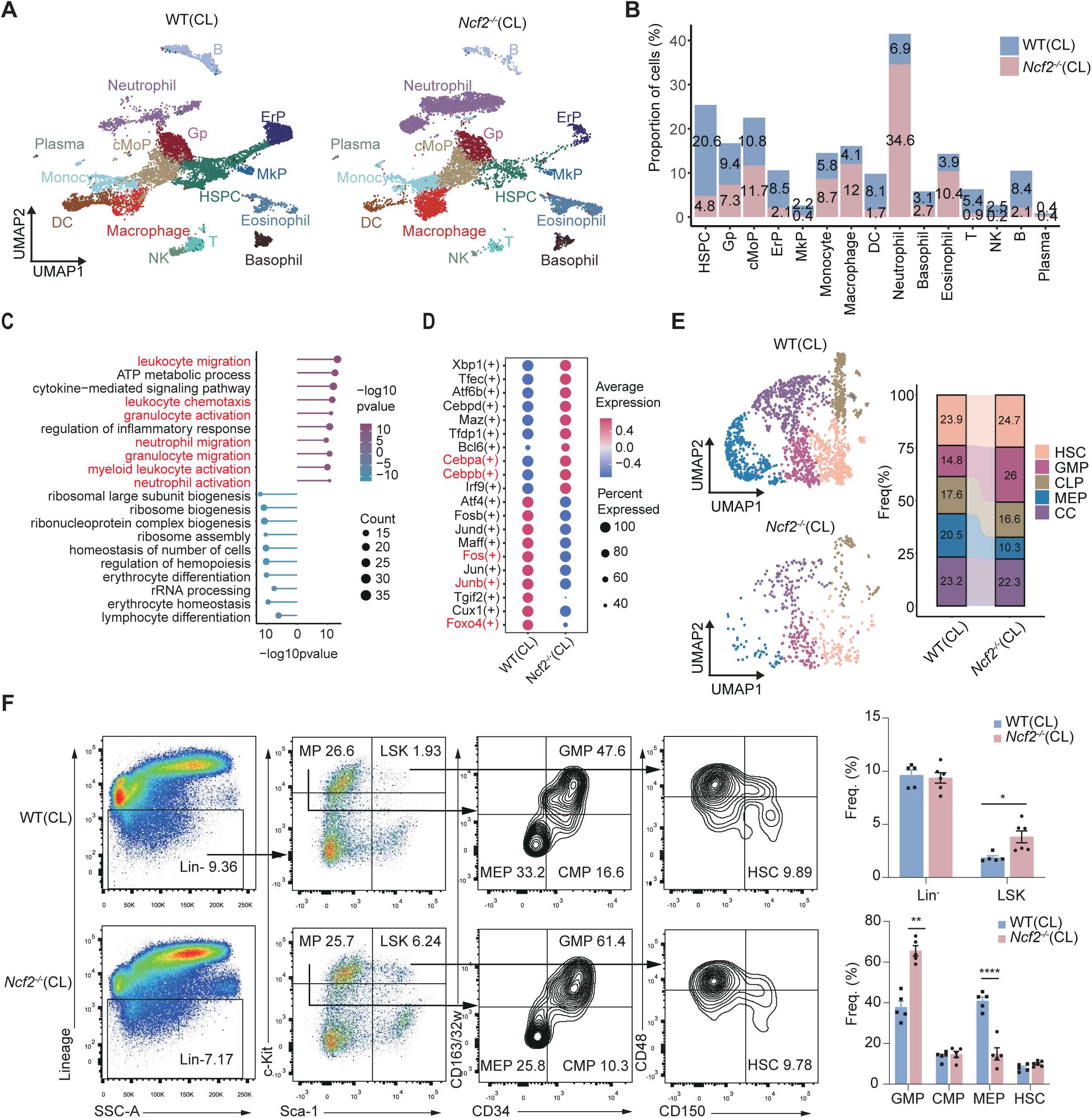
Systemic inflammation leads to hematopoietic disturbances in the BM of CGD mice **(A)** UMAP of hematopoietic and immune cells from CL WT (n=9687) and CL *Ncf2^-/-^* (n=9528) mice BM. **(B)** Stacking bar plots illustrating the frequency of each cell type shown in (A). **(C)** Lollipop chart of enrichment analysis for differentially expressed genes in the HSPC subgroups (CL *Ncf2^-/-^* vs CL WT), with red bars representing upregulated gene items and green bars representing downregulated items (Top 10). **(D)** Transcription factors analysis in the HSPC across two groups conducted by SCIENIC. **(E)** UMAP plots displaying sub-clusters of HSPC in CL WT and CL *Ncf2^-/-^* mice. **(F)** Representative scatter plots and contour plots revealing hematopoietic stem and progenitor cell compartments in bone marrow of CL WT and CL *Ncf2^-/-^* mice. Bar chart on the right depicting the proportion of cells. HSPC: Hematopoietic Stem and Progenitor Cells, HSC: Hematopoietic Stem Cells, MEP: Megakaryocyte/Erythroid Progenitors, GMP: Granulocyte/Monocyte Progenitors, CMP: Common Myeloid Progenitors, LSK: Lin^-^Sca-1^+^ c-Kit^+^ cells.

Analysis of differentially expressed genes (DEGs) showed that inflammatory factors *S100a8* and *S100a9* were the most significantly up-regulated transcripts in the BM cells of CL *Ncf2*^-/-^ mice (**Figure S11B**). Additionally, there was an up-regulation of myeloid characteristic genes, such as *Ngp* (Neutrophilic granule protein), *Lyz2* (lysozyme 2), *Camp* (cathelicidin anti-microbial peptide), and *Retnlg* (resistin like gamma) (**Figure S11B**). Subsequent GO enrichment analysis in HSPC subsets indicated that signaling pathways involved in leukocyte migration, chemotaxis, neutrophil activation, and inflammatory response were significantly up-regulated in the CL *Ncf2^-/-^* group (**Figure 7C**). While, the down-regulated pathways were predominantly related to ribosomal functions, lymphocytes and erythrocytes differentiation (**Figure 7C**). By utilizing the SCENIC transcription factor (TF) analysis, we further elucidated key regulatory molecules potentially driving HSPC differentiation under inflammatory conditions. In CL *Ncf2^-/-^* HSPCs, myeloid differentiation-related TFs such as *Cebpa* and *Cebpb*, exhibited enhanced activity. Conversely, the transcriptional activity of TFs such as *Fos, Junb*, and *Foxo4*, which were crucial for hematopoietic homeostasis were decreased (**Figure 7D**). Additionally, bulk RNA-seq of BM Lin^-^ cells from both groups revealed a significant upregulation of pro-inflammatory cytokine genes such as *Il1b, Il6,* and *Tnf* in CL *Ncf2^-/-^* Lin^-^ cells (**Figure S11C**). Due to the heterogeneity of HSPCs, these cells were further divided into 5 subsets based on specific gene expression, including hematopoietic stem cells (HSC), Granulocyte-monocyte progenitor cells (GMP), Common Lymphoid Progenitor cells (CLP), Megakaryocyte-erythrocyte progenitor cells (MEP) and cycling cells (CC) (**Figure 7E**). There was a notable increase in the proportion of GMP cells, while the proportions of CLP and MEP decreased (**Figure 7E**). Subsequently, flow cytometry analysis was conducted to corroborate the findings from the scRNA-seq profiles. The results indicated that in the BM of CL *Ncf2^-/-^* mice, the proportions of Lin^-^Sca-1^+^c-Kit^+^cells (LSK), CMP, and GMP were significantly increased, whereas the proportion of MEP was substantially reduced, with the HSC population remaining constant (**Figure 7F**).

In summary, as revealed by scRNA-seq profiling and flow cytometry within the CGD mouse model, the activation of hematopoietic precursor cells by multiple pro-inflammatory cytokines such as *Il1b, Il6,* and *Tnf* triggered emergency hematopoiesis. Driven by a sophisticated transcriptional regulatory network, the hematopoietic cell compartment in CGD mice underwent significant remodeling, characterized by a pronounced myeloid skew and the suppression of non-myeloid hematopoiesis. We also profiled the cellular alterations in gut tissues from both CL WT and CL *Ncf2^-/-^* mice. In contrast to the dramatic alteration in the BM cells, we observed minimal alterations in the gut tissue (**Figure S12A and S12B**).

### *Morrbid* genetic deficiency reduced excessive inflammatory response in CGD mice by modulating myeloid apoptosis

The excessive activation and proliferation of myeloid cells were critical factors that contributed to persistent inflammatory responses and tissue damage. Multiple studies have shown that polymorphic neutrophils (PMNs) in CGD exhibited an apoptotic defect, which prolonged the resolution of inflammation.^35–38^ Through bulk RNA-seq analysis, we identified that the upregulated DEGs in Lin^-^ cells of CL *Ncf2^-/-^* mice were significantly enriched in pathways associated with apoptosis inhibition (**Figure S13A**). Additionally, the pro-survival gene *Morrbid* was upregulated (**Figure 8A**).

**Figure 8.**
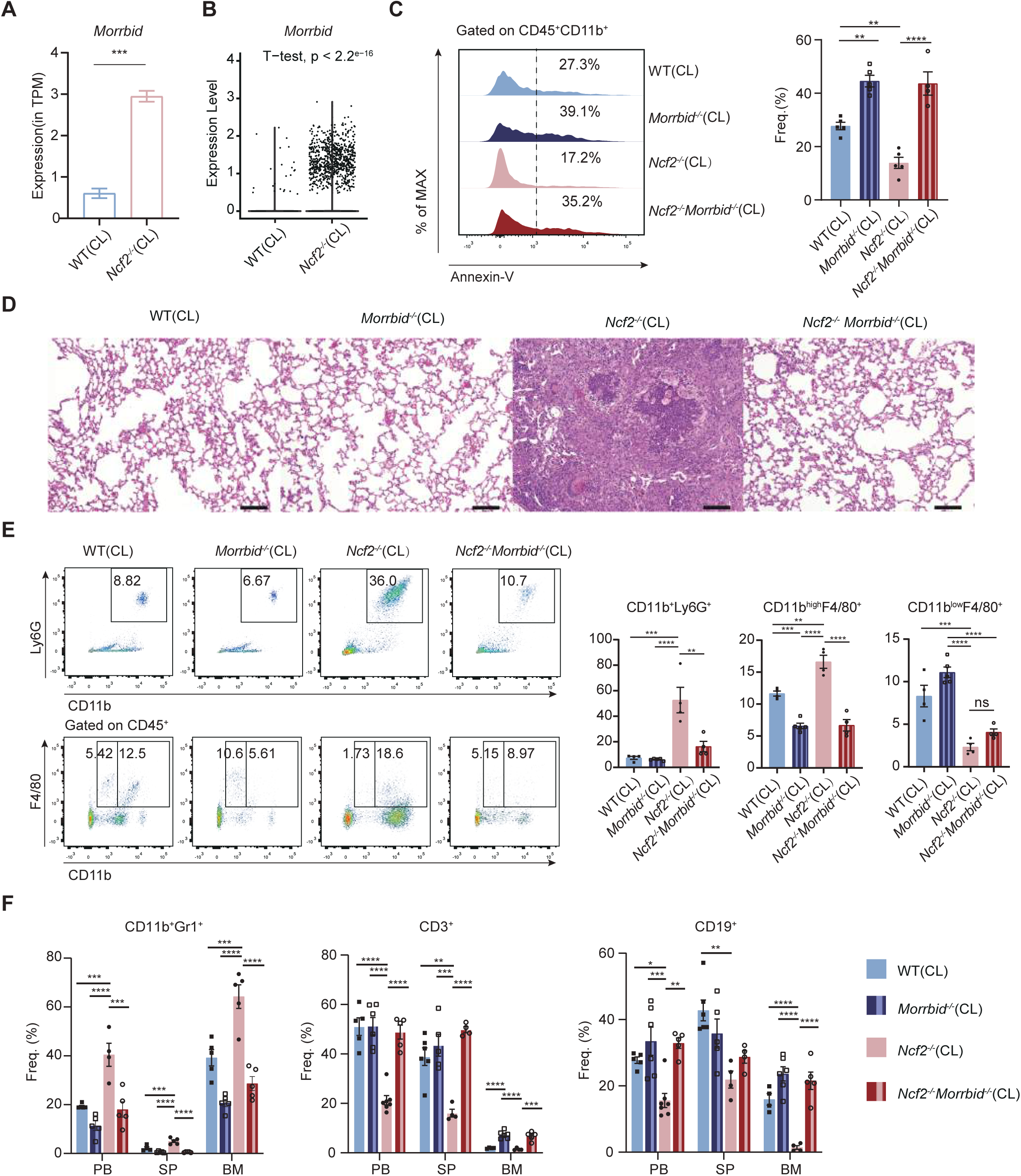
Genetic deficiency in *Morrbid* alleviates hyperinflammation in CGD mice **(A)** Expression of the pro-survival gene *Morrbid* in Lin^-^cells (Bulk RNA-seq) . **(B)** Expression of the pro-survival gene *Morrbid* in neutrophils of mice BM (scRNA-seq). **(C)** Apoptosis level of myeloid cells from CL WT, CL *Morrbid^-/-^*, CL *Ncf2^-/-^*, and CL *Ncf2^-/-^Morrbid^-/-^* mice analysed by Annexin-V flow cytometry. **(D)** HE staining of lung tissues from CL WT, CL *Morrbid^-/-^*, CL *Ncf2^-/-^*, and CL *Ncf2^-/-^Morrbid^-/-^* mice. Scale bar: 100μm. **(E)** Representative flow cytometry scatter plots of neutrophils and macrophages from lung tissues of CL WT, CL *Morrbid*^-/-^, CL *Ncf2*^-/-^, and CL *Ncf2*^-/-^*Morrbid*^-/^- mice. Bar plots displaying the proportions of CD11b^+^Ly6G^+^ neutrophils, CD11b^high^F4/80^+^ MDMs, and CD11b^low^F4/80^+^ resident macrophages. **(F)** Proportions of granulocytes and lymphoid cells in the peripheral blood (PB), spleen (SP), and bone marrow (BM) of CL WT, CL *Morrbid*^-/-^, CL *Ncf2*^-/-^, and CL *Ncf2^-/-^Morrbid^-/-^* mice analyzed by flow cytometry.

Furthermore, in the scRNA-seq dataset, the upregulated DEGs in CL *Ncf2^-/-^* mice’s neutrophils were notably concentrated in apoptosis inhibition pathways, and *Morrbid* gene expression was elevated (**Figure 8B** and **Figure S13B**). *Morrbid* is a lncRNA specifically expressed in short-lived myeloid cells (such as neutrophils, eosinophils, and monocytes) and their progenitor cells, playing a crucial role in inhibiting the apoptotic process.^12^ Our previous studies demonstrated important pathophysiological roles of *Morrbid* in leukemogenesis and clonal hematopoiesis.^39–41^ We hypothesize that genetic deficiencies in *Morrbid* directly regulate apoptosis in myeloid cells of the CGD mice and alleviate associated inflammatory responses. By crossbreeding *Morrbid^-/-^* mice with *Ncf2^-/-^* mice, we constructed a *Morrbid^-/-^Ncf2^-/-^*double knockout model to explore the impact of *Morrbid* genetic deficiencies on systemic inflammation in a CGD context. Flow cytometry showed that CL *Ncf2^-/-^* mice had reduced apoptosis levels of CD11b^+^ mature myeloid cells compared with CL WT mice, while *Morrbid^-/-^*genetic deficiency significantly promoted the apoptotic process (**Figure 8C**).

To further explore the effects of *Morrbid* deficiency on pulmonary inflammation in CGD mice, we performed H&E staining of the lung tissues from CL WT, CL *Morrbid^-/-^*, CL *Ncf2^-/-^*, and CL *Ncf2^-/-^Morrbid^-/-^* mice. Compared to CL *Ncf2^-/-^*, there was a marked reduction in inflammatory cell infiltration and granuloma formation in the lung tissue of CL *Ncf2^-/-^Morrbid^-/-^* mice, with no significant consolidation observed (**Figure 8D**). Flow cytometry results showed that the proportion of CD11b^high^F4/80^+^ MDMs in the lung tissues of CL *Morrbid^-/-^* mice was significantly lower than that of CL WT mice, while the proportion of CD11b^low^F4/80^+^ resident macrophages was increased (**Figure 8E**). Additionally, the CL *Ncf2^-/-^Morrbid^-/-^* group showed a significant reduction in Ly6G^+^ neutrophils and CD11b^high^F4/80^+^ MDMs compared to the CL *Ncf2^-/-^* group (**Figure 8E**). Therefore, we demonstrated that *Morrbid* genetic deficiency effectively reduced inflammatory cell infiltration in the lung tissues of CGD mice. Flow cytometry analysis was utilized to further investigate the impact of *Morrbid* genetic deficiency on hematopoiesis. As shown in **Figure S13C**, the results indicated that, compared to that in CL WT, *Morrbid* deficiency significantly reduced the proportion of CMP, aligning with previous findings.^12^ Compared to CL *Ncf2^-/-^*, the proportion of CMP and GMP were significantly decreased in CL *Ncf2^-/-^Morrbid^-/-^*mice, while the proportion of MEP was restored (**Figure S13C**). Subsequent analysis of mature immune cell fractions revealed that the proportion of CD11b^+^Gr1^+^ granulocytes in the PB, SP, and BM of CL *Ncf2^-/-^Morrbid^-/-^* mice was significantly decreased than that in CL *Ncf2^-/-^* Mice (**Figure 8F** and **Figure S13D**). In contrast, the proportions of CD3^+^ T lymphocytes and CD19^+^ B lymphocytes in the PB, SP, and BM increased significantly, suggesting a recovery of adaptive immune function (**Figure 8F**). In summary, our results solidly demonstrated that along with targeting MIF pathway or IL1R1 pathway, *Morrbid* deficiency also alleviate abnormal myeloid hematopoiesis and disease progression in the natural CGD mice.

## DISCUSSION

Through different environmental exposures (SPF or CL), we fully explored the impact of environmental pathogens on the phenotype of mice with a CGD genetic background. The CL environment contains certain pathogens sensitive to the *Ncf2*^-/-^ immune-deficient mice, as evidenced by our microbiota studies. WT mice are able to eliminate such pathogens under CL conditions, whereas *Ncf2*^-/-^ mice raised in this environment recaptured the natural history of CGD patients. With the assistance from regular single-cell and spatial transcriptomic studies and verification with other experimental approaches, we decoded several important events induced by the immune aberrations caused by *Ncf2* loss. Importantly, we developed three different but very possibly correlated perturbations to mitigate CGD progression: targeting *Il1r1*, *Mif* or *Morrbid*. Such perturbations are very likely applicable for controlling systemic inflammation and granuloma formation in CGD.

Patients with CGD are perpetually at risk of opportunistic infections from catalase-positive pathogens such as Aspergillus, Staphylococcus, Klebsiella, and Burkholderia. However, manifestations of infection are infrequently observed in NOX2-deficient mouse models at the SPF-grade experimental facility. This discrepancy is very likely attributed to environmental variables, given that these mice are maintained in pathogen-free conditions, which starkly contrast the intricate microbial landscapes encountered in natural settings.^42^ We speculated that exposing Nox2-deficient mice to an environment with a certain fungal and bacterial abundance will simulate the disease state as same as that in CGD patients. In our research, C57BL/6 WT mice and *Ncf2*^-/-^ mice were placed in both SPF and CL environments to assess the impact of environmental exposure. Consistent with prior findings, no signs of infection were detected in *Ncf2*^-/-^ mice in the SPF setting.^43^ However, CL *Ncf2*^-/-^ mice exhibited pronounced pulmonary infections and spontaneous necrotizing granulomatous lesions. Notably, the pathogenic bacteria of lung infections commonly seen in CGD patients, such as Klebsiella and Staphylococcus, were the primary bacterial flora in the lungs of CL *Ncf2*^-/-^ mice, while almost absent in the other three scenarios.^44,45^ These results support the hypothesis that a granulomatous model for CGD can be developed solely through environmental manipulation. Nevertheless, ITS bacterial sequencing revealed that the Talaromyces was the predominant fungal group in the lung tissue of CL *Ncf2*^-/-^ mice. Although this fungus is prevalent among patients with immunodeficiencies like HIV, it has yet to be reported in CGD cases, possibly because laboratory conditions still unable to fully replicate the complex environments of daily human life.

Neutrophils, as pivotal effector cells of innate immunity, are rapidly recruited to inflammatory sites guided by chemokines, lipid chemo-attractants, complement factors, formyl peptides, and their respective receptors.^46^ At these sites, neutrophils eradicate pathogens through mechanisms such as phagocytosis, degranulation, and the release of ROS and neutrophil extracellular traps (NETs).^46^ The deficiency in NOX2 triggers the production of additional inflammatory mediators, resulting in the excessive accumulation of neutrophils and unresolved inflammation. In CL *Ncf2*^-/-^ mice, neutrophils are the predominant cell type within the immune microenvironment of granulomas and are strategically located in the core areas, underscoring their central role in the development of CGD-associated granulomas. Within these cores, we identified a subgroup of neutrophils characterized by high expression of NOS2, which displays pro-inflammatory transcriptional profiles. NOS2, in the presence of inducing stimuli and the substrate L-arginine, can produce significant amounts of NO, which has extensive bactericidal activity.^47^ Therefore, the elevated expression of NOS2 in neutrophils within CGD granulomas likely acts as a compensatory response to the deficiency of ROS in phagocytes, thus enhancing host defense to some extent. Consistent with this, the production of intracellular NO in neutrophils from CGD patients increases in a time-dependent manner upon stimulation with lipopolysaccharide (LPS) and a calcium ionophore.^48^ Tsuji *et al* has demonstrated that PMNs from CGD patients showed increased production of NO following phagocytosis.^49^ Furthermore, a study by Ahlin *et al* indicated that interferon-γ treatment enhanced NO production in PMNs of CGD patients to enhance bactericidal capacity.^50^

The pulmonary macrophage population exhibits significant diversity and complexity, and are classified into alveolar macrophages, interstitial macrophages, and MDMs based on surface markers, location, and origin. Alveolar and interstitial macrophages, as resident pulmonary macrophage populations, play a crucial role in defending against pathogens and clearing cellular debris, thereby maintaining pulmonary homeostasis.^51^ However, severe infections can lead to a substantial depletion of alveolar macrophages, a phenomenon known as the "macrophage disappearance reaction".^51^ When the remaining resident macrophages cannot be replenished in the short term, the recruitment of monocytes is often essential for restoring their numbers.^52^ During this process, the depletion of resident macrophages significantly diminishes survival rates and intensifies lung injury.^53^ In lung tissues of CGD mice, infection leads to notable depletion of resident macrophages, with a significant increase in monocyte-derived macrophages (Mac1), which become the predominant macrophage type in the granulomatous regions. These MDMs exhibit high reactivity and plasticity, and are "polarized" by cytokines, dead cells, and microbial products.^54^ Arginine metabolism is crucial in this context, influencing the polarization of M1 and M2 macrophages through the activation of the NOS2 and arginase pathways, respectively.^55^ The NOS2 pathway promotes M1 macrophage differentiation, exerting cytotoxic effects through the production of citrulline and NO to manage bacterial infections. Conversely, the arginase pathway enhances the production of proline and urea, driving M2 polarization to promote wound healing. In the peripheral regions of CGD granulomas, we identified a subgroup of MDMs highly expressing NOS2 and ARG1. This indicates that the formation of necrotic granulomas associated with CGD may require a combination of pro-inflammatory macrophages with bactericidal activity and anti-inflammatory macrophages that promote healing to modulate the immunopathological response. Additionally, macrophages at the edge of granulomas also highly express *Timp2*, *Mmp9*, *Mmp12*, and *Mmp14*, suggesting their pro-fibrotic characteristics.^56^

The excessive inflammatory response leading to abnormal accumulation of neutrophils and MDMs is fundamental in the formation of pulmonary granulomas in CGD, with the upregulation of pro-inflammatory mediators serving as a key driver. Previous studies have confirmed the critical roles of IL-1α and IL-1β in the inflammatory responses of various CGD disease models, including serum-induced sterile arthritis,^57^ sodium periodate-induced sterile peritonitis,^58^ zymosan-induced sterile pneumonia,^18^ and Aspergillus-induced fungal pneumonia^17^. IL-1α and IL-1β function by binding to the widely expressed IL-1 receptor (IL-1R), thereby initiating the production of downstream chemokines and cytokines. In our *Ncf2^-/-^Il1r1^-/-^* DKO mice, systemic inflammation was effectively controlled, underscoring the role of essential signaling molecules in CGD’s hyperinflammatory condition. Investigating mediators that may influence CGD-related inflammatory states is crucial for developing precise immunomodulatory strategies. In scRNA-seq and spatial transcriptomic datasets, we noted a significant upregulation of *Mif* in lung neutrophils and macrophages of CL *Ncf2*^-/-^ mice, particularly in specific Neu6 and Mac1 subsets. We proposed that MIF plays a critical role in driving the inflammatory progression of CGD. Pharmacological inhibition of MIF markedly reduced neutrophil infiltration in CGD lung tissue and corrected myeloid hematopoiesis bias. Additionally, pseudo-time analysis indicated similar expression patterns for *Mif* and *Il1b* in neutrophils and macrophages, suggesting a regulatory relationship. Research has shown that MIF facilitated the interaction between NLRP3 and the intermediate filament protein vimentin, thus mediating the activation of the NLRP3 inflammasome.^32^ A study by Wang *et al* demonstrated that macrophages enhance the progression of oral squamous cell carcinoma through the MIF/NLRP3/IL-1β axis.^59^ In systemic lupus erythematosus, snRNP immune complexes induce monocytes to produce MIF and mediate IL-1β production through NLRP3 inflammasome activation.^60^ Through ELISA and western blot assays, we have confirmed the critical role of the MIF/NLRP3/IL-1β signaling pathway in promoting the inflammatory response in CGD.

In our study, the significant expansion of myeloid cells, particularly neutrophils and MDMs, constitutes the pathological basis for the formation of CGD granulomas. This uncontrolled proliferation of cells may result from a myeloid bias triggered by systemic inflammatory stimuli, as well as delayed apoptosis in myeloid cells. *Morrbid*, a lncRNA specifically expressed in myeloid cells and their precursors.^12^ However, sustained expression of *Morrbid* may lead to prolonged survival of these cells, which is associated with the development of various chronic inflammatory diseases and hematologic disorders.^39–41^ We observed upregulation of *Morrbid* in the BM neutrophils and Lin^-^ cells of CGD mice. Knocking out the *Morrbid* significantly enhanced apoptosis in myeloid cells of CGD mice and reduced the accumulation of neutrophils in the BM and peripheral organs, particularly in lung tissues, thereby effectively suppressing the inflammatory response. Our results emphasized the crucial role of *Morrbid* in regulating neutrophil apoptosis in CGD mice and proposed novel therapeutic approaches for mitigating the associated hyperinflammation.

In summary, the results of our study are encapsulated in a schematic diagram (**Figure S14**). Initially, by manipulating the living conditions of mice with a CGD genetic background, we successfully mimicked the typical pulmonary infections observed in patients and formed the characteristic necrotic granulomatous lesions. These manifestations of infection are likely driven by prevalent CGD pathogens such as Staphylococcus and Klebsiella, along with the opportunistic fungus Talaromyces. Through scRNA-seq analysis, spatial transcriptomic analysis, and flow cytometry, we identified neutrophils and MDMs as the primary immune constituents of the CGD pulmonary granulomas. Spatially, the NOS2^high^ neutrophil subpopulation with pro-inflammatory transcriptional characteristics is located in the granuloma core area, while the specific macrophage subpopulation marked by NOS2 and ARG1 is mainly distributed in the granuloma margin region, and facilitating the formation of fibrotic encapsulation through the expression of EMC remodeling genes. Mechanistically, we propose that the excessive inflammatory response leading to myeloid cell accumulation is a fundamental condition for the formation of granulomas. 4IPP treatment can inhibit inflammatory cell infiltration in the lungs of CGD mice by regulating the MIF/NLRP3/IL1β signaling pathway, thereby reducing the myeloid bias caused by emergency hematopoiesis. Additionally, the genetic deficiency of *Morrbid* can effectively limit inflammation and granuloma formation by promoting the specific apoptosis of myeloid cells.

## CONCLUSION

In this study, we generated a mouse model for natural CGD with deficiency in *Ncf2* gene but without additional pathogen challenge. We analyzed the pathological phenotypes and possible pathogens in the clean conditon, decoded immune events by regular single-cell RNA-sequencing and spatial transcriptomics, and developed the three different perturbations. Our research delves deeply into the mechanisms underlying granuloma formation and paves the way for novel interventions to suppress hyperinflammation in CGD.

## Supporting information

### Acknowledgements

We thank our colleagues for technical support, critically reading our manuscript, and their suggestions to improve the manuscript. We would also like to thank Drs. Zheng and Yu (Tianjin Medical University School of Basic Medicine), and Cheng (Chinese Academy of Medical Sciences, National Clinical Research Center for Blood Diseases) for sharing mice.

## METHODS

### Animals

C57BL/6 WT, *Morrbid^+/-^, Il1R1^+/-^*, and *Ncf2^+/-^* mice were purchased from Cyagen Biosciences. *Morrbid^+/-^, Il1R1^+/-^,* and *Ncf2^+/-^* mice were bred to obtain homozygous mice through self-crossing. *Morrbid^-/-^* and *Il1R1^-/-^*mice were then crossbred with *Ncf2^-/-^* mice for two generations to generate *Morrbid^-/-^Ncf2^-/-^* and *Il1R1^-/-^Ncf2^-/-^* mice. Depending on experimental requirements, mice were housed in SPF or CL grade environments. Mice designated for CL environment exposure were initially raised under SPF conditions and subsequently transferred to the CL conditions for a one-month exposure. All experimental mice were approximately 8 weeks old.

### Bronchoalveolar lavage (BAL)

To collect BAL samples, we performed three consecutive lavages with 1 ml of pre-cooled 3% fetal bovine serum (FBS) phosphate-buffered saline (PBS). Subsequently, the BAL samples were centrifuged at high speed to obtain the supernatant, which was then used for ELISA.

### Preparation of cell suspensions

Fresh mouse lung tissues were minced and then digested at 37°C for 45 minutes with 1 mg/ml type IV collagenase. The resulting suspension was passed through a 40 μm cell strainer. BM cells were harvested from the femurs and tibias, and filtered through a 40 μm cell strainer. Mouse spleens were crushed and similarly filtered through a 40 μm cell strainer. The filtrates obtained were treated with red blood cell lysis solution, then washed and resuspended for further use.

### Isolation of Lin-negative BM cells

According to the manufacturer’s instructions, Lin^-^ cell populations were isolated from mouse BM samples using negative selection with the EasySep™ Mouse Hematopoietic Progenitor Cell Isolation Kit (STEMCELL, 19856A).

### Histology

Fresh lung tissues were fixed in 4% paraformaldehyde for 24 hours, followed by embedding in paraffin. Sections of 4 μm thickness were stained with hematoxylin and eosin (H&E) or Masson’s trichrome. The slides were then imaged under an optical microscope (Olympus, Japan).

### Enzyme-linked immunosorbent assay (ELISA)

The Mouse MIF (Macrophage Migration Inhibitory Factor) ELISA Kit (Elabscience, E-EL-M0771) and the Mouse IL-1β ELISA Kit (MULTI SCIENCES, EK201B/3-48)

were used to measure the concentrations of MIF and IL-1β proteins, respectively, following the manufacturer’s instructions. The protein concentrations of IL-6, IFN-γ, and TNF-α in serum were determined using the Proinflammatory Panel 1 (mouse) kits (MSD, K15048D).

### Western blot analysis

Proteins (30 µg) were extracted from lung tissues and separated by 10% SDS-polyacrylamide gel electrophoresis. The separated proteins were then transferred onto 0.45 µm PVDF membranes. The membranes were blocked with a protein-free rapid blocking solution (EpiZyme, PS108) at room temperature for 30 minutes. They were then incubated overnight at 4°C with antibodies against NLRP3 (1:1000, Abcam, AB270449) and β-Actin (1:1000, ZSGB-BIO, TA-09). Subsequently, the membranes were incubated for 1 hour at room temperature with HRP-labeled Goat Anti- Rabbit lgG (H+L) (1:1000, EpiZyme, LF102) or Goat Anti-Mouse IgG/HRP (1:1000, Solarbio, SE131). Protein bands were detected using StarSignal Chemiluminescent Assay Kit (Genstar, E171-1) and visualized using a digital imaging system (BioRad). All samples were normalized to Actin and analyzed using ImageJ software(version 1.53t).

### Flow cytometric analysis

Single cells isolated from mouse BM, PB, SP, and lung tissues were resuspended in PBS. For cell surface protein staining, the cells were incubated with conjugated antibodies (1:100) at 4°C for 30 minutes. For intracellular proteins, the True-Nuclear™ Transcription Factor Buffer Set (BioLegend, 424401) was used prior to antibody staining. The antibodies used included CD45.2-PerCP/Cyanine 5.5 (BioLegend, 109828), CD11b-Pacific Blue (BioLegend, 101224), CD11b-PE (BioLegend, 101208), CD11b-FITC (BioLegend, 101206), Ly6G-PE (BioLegend, 127608), F4/80-APC/Cyanine7 (BioLegend, 123118), Arg1-Alexa Fluor^TM^700 (eBioscience, 56-3697-82), iNOS-APC (eBioscience, 17-5920-82), cKit-APC (BioLegend, 105812), Sca-1-APC/Cyanine7 (BioLegend, 108126), CD48-PE/Cyanine7 (BioLegend, 103424), CD150-PE/Cyanine5 (BioLegend, 115912), CD16/32-PE/Cyanine7 (BioLegend, 101318), CD34-FITC (eBioscience, 11-0341-85), Gr1-PE (BioLegend, 108408), Gr1-Brilliant Violet 421 (BioLegend, 108434), CD19-APC (BioLegend, 152410), CD3-PE (BioLegend, 100206), TER-119-PE (BioLegend, 116208), B220-PE

(BioLegend, 103208) and Annexin V-PE (KeyGEN, KGA1018). After washing away unbound antibodies, the cells were analyzed using a BD LSRFortessa (BD Biosciences, USA) and FlowJo software (version 10.0.7). The Reactive Oxygen Species Assay Kit (Beyotime, S0033S) with the fluorescent probe DCFH-DA was used for the reactive oxygen species assay, following the manufacturer’s instructions.

### scRNA-seq and data processing

The lung tissue single cells were resuspended in RPMI 1640 culture medium containing 4% fetal bovine serum after dissociation. Cell viability and counting were performed using AO/PI staining with the Countstar® Rigel S2 Fluorescent Cell Analyzer, and the cell concentration was adjusted to 1×10^6^ cells per milliliter. Subsequently, the SeekOne® Digital Droplet Single Cell 3’ Library Preparation Kit (SeekGene, No.K00202) was used to prepare the single-cell RNA-Seq library. The library preparation process included generating emulsion gel beads (GEMs), barcode tagging, GEM-RT Clup, cDNA amplification and quantification, and library construction, all executed according to the manufacturer’s protocols. The prepared sequencing libraries were purified using SPRI beads and quantified using KAPA Library Quantification Kits (Roche, KK4824) to ensure library quality. Sequencing was performed on the Illumina NovaSeq 6000 platform, using a 150-base-pair paired-end reads format. After sequencing, raw data were processed with the SeekSpace tools (v1.0.0, Seekgene) to align reads to the mouse reference genome (mm10) and generate a gene-cell expression matrix for subsequent analysis.

The single-cell expression matrix and sample information from SPF WT, SPF *Ncf2^-/-^*, CL WT, and CL *Ncf2^-/-^* lung tissues were integrated using the R package Seurat (v4.2.0) to facilitate subsequent comprehensive analysis. Cells with more than 50,000 transcripts, over 6,000 genes, mitochondrial gene expression greater than 5%, or ribosomal gene expression exceeding 50% were filtered out. Following the stringent quality control, 11305 cells in the SPF WT group, 7948 cells in the SPF *Ncf2^-/-^* group, 7912 cells in the CL WT group, and 8131 cells in the CL *Ncf2^-/-^*group were retained for further analysis. Gene expression data were normalized using the default parameters of the NormalizeData function, and highly variable genes were identified by the FindVariableFeatures function. The top 2,000 highly variable genes were scaled using the "ScaleData" function. Dimensionality reduction was performed using Principal Component Analysis (PCA), with the top 50 principal components selected for clustering analysis. To eliminate potential batch effects between different groups, we utilized the RunHarmony function for correction. A resolution of 0.5 was used for unbiased clustering, and UMAP was employed to visualize the dataset on a two-dimensional plane. Using the FindAllMarkers function, signature genes for each cluster were identified and compared with known lineage-specific genes, enabling the annotation of cell types. In addition, the FindMarkers function was used to analyze differential expression between any two groups, thereby generating a list of DEGs. GO enrichment analysis was conducted on these DEGs, setting a P-value of less than 0.05 as statistically significant enrichment. Concurrently, scRNA-seq analyses were also performed on BM and colon tissues from both CL WT and CL *Ncf2^-/-^* groups. The data processing workflow for these tissues was consistent with that used for lung to ensure methodological consistency.

### Focused analysis of specific cell clusters

In the lung single-cell dataset, the subset function within the Seurat package was utilized to selectively extract neutrophils and mononuclear macrophages for in-depth analysis. Resolution parameters of 0.4 for neutrophils and 0.2 for mononuclear macrophages were applied, facilitating unbiased cluster analysis to explore the subpopulation structure. For the BM dataset, a resolution of 0.3 was used to further examine the HSPC population. These steps followed the same analysis procedures previously applied to the entire dataset, including data normalization, selection of highly variable genes, data scaling, PCA dimensionality reduction, and UMAP visualization.

In single-cell trajectory analysis, we utilized Monocle 3 (version 1.0.0) to explore the developmental trajectories of neutrophil and mononuclear/macrophage compartments. Initially, the scRNA-seq dataset was normalized, standardized, and clustered using Seurat, after which the data was constructed into Monocle3 objects for further analysis. By applying the learn_graph function, we conducted pseudotime trajectory analysis on these cells. Within the neutrophil population, the Neu1 expressing the immature neutrophil genes was selected as the root state of the trajectory. For the mononuclear/macrophage population, the Mon1 cell population expressing the typical monocyte markers was selected as the starting point. After constructing the developmental trajectories, the plot_cells function was used for visualization. Additionally, the plot_genes_in_pseudotime function was employed to display the dynamic trends in the expression of specific genes over pseudotime.

To reconstruct the gene regulatory network, we used SCENIC software (version 0.11.2) to identify transcriptional regulatory factors in HSPC subpopulations and assess the transcriptional activity of these regulators across different groups.

### Spatial transcriptomics

First, the lung tissue samples from CL WT and CL *Ncf2*^-/-^ were embedded in the optimum cutting temperature (OCT) compound (SAKURA,Cat#: 4583). After cutting these samples into 10-micron thick slices, RNA was extracted, and its quality was assessed to ensure that the RNA Integrity Number (RIN) > 7. Before executing the complete spatial gene expression library construction process, tissue optimization experiments were necessary to determine the optimal permeabilization time. Once the permeabilization time is established, the tissue samples were subjected to the corresponding permeation treatment, releasing intracellular mRNA to bind with capture probes and undergo reverse transcription into cDNA. Subsequent steps include double-stranded cDNA synthesis, denaturation recovery, amplification, and purification to obtain sufficient quantities of complete cDNA. Subsequently, the purified cDNA samples were used for library construction (SeekSpace sc-Spatial Labeling Kit V 1.0 or BMKMANU S1000 Gene Expression Kit). The detailed steps included fragmentation, end repair, adapter ligation, bead purification, and PCR quantification. After library construction, the concentration was measured by Qubit assays (Thermo, Q32854) and appropriately diluted as needed. The size of the library

fragments was checked using the 2100 Bioanalyzer Instrument, ensuring that the fragment lengths typically ranged between 300 and 700 bp. The qualified libraries were then sequenced on an Illumina NovaSeq 6000.

### Preprocessing and analysis of spatial transcriptomics data

Raw base calls from SeekSpace were demultiplexed and converted to FASTQ format using the bcl2fastq2 (version 2.2.0, Illumina). The FASTQ files were aligned to the mouse reference genome assembly mm10 and a count matrix was generated using SeekSpace tools (version 1.0.0, Seekgene).While, the upstream analysis of the raw data from BMKMANU S1000 was executed using BSTMatrix (version 1.0).

The spatial transcriptomics data, including expression matrices, sample information, and spatial coordinates, were loaded using the CreateSeuratObject function to construct a Seurat object. Spatial spots with high expression of erythrocyte-associated genes were filtered out (>2%). Subsequently, data normalization was performed using the LogNormalize algorithm. The FindVariableFeatures function was used to identify highly variable genes (top 2000), and expression data for these genes were scaled using the ScaleData function. Principal component analysis was conducted using the RunPCA function with default parameters. Unsupervised clustering was performed using FindNeighbors (dims = 1:30) and FindClusters functions (resolution = 1.5). Data dimensionality reduction and visualization were accomplished using the Uniform Manifold Approximation and Projection (UMAP) method.

Using the Robust Cell Type Decomposition (RCTD) algorithm, we inferred the cellular composition of spatial spots for CL *Ncf2^-/-^*and CL WT samples based on annotated information identified from our scRNA-seq transcriptome profiles. To accurately annotate the spots, cell types with fewer than 50 cells in the single-cell dataset were filtered out.

### RNA-sequencing and analysis

RNA was prepared from BM Lin^-^cell suspension sorted by magnetic beads. Total RNA was isolated using the QIAzol Lysis Reagent (Qiagen, 79306), with all samples displaying an RIN greater than 6.5. The RNA-seq library was then prepared using the Illumina Stranded mRNA Prep, Ligation kit and sequenced on the Illumina NovaSeq 6000 platform.

During the analysis of transcriptome sequencing data, we initially utilized the TrimGalore software (version 0.6.7) to preprocess the raw sequencing data, removing low-quality reads and adapter sequences. Subsequently, the processed data underwent quality control using FastQC (version 0.11.9). The sequencing fragments were then mapped to the mouse reference genome using the Hisat2 (version 2.2.0) alignment tool. The featureCounts tool (version 2.0.1) was used to efficiently count the reads mapped to exons, thereby constructing the raw count matrix needed for subsequent analysis. The raw count matrix was loaded into R version 4.2.0, and gene reads were normalized using TPM (Transcripts Per Million) for further analysis.

### 16s rRNA/ITS sequencing

The lung tissues from four groups (SPF WT, SPF *Ncf2^-/-^*, CL WT, CL *Ncf2^-/-^*, each group n=3) were homogenized, and total DNA was extracted using the FastPure Soil DNA Isolation Kit (Magnetic bead)(MJYH, shanghai, China). The NEXTFLEX Rapid DNA-Seq Kit (Bioo,5144-08) was used for library construction, followed by sequencing on Nextseq 2000 platform. Raw sequencing data were quality-controlled with fastp (version 0.20.0) and assembled using FLASH (version 1.2.11). Sequences were clustered into OTUs at 97% similarity using UPARSE software (version 11). All downstream data analyses were performed on the Majorbio Cloud Platform (https://cloud.majorbio.com). Specifically, alpha diversity indices, including Chao and Shannon indices, were calculated using mothur software (version 1.30.2) to assess species richness and diversity. Principal Coordinates Analysis (PCoA) based on the Bray-Curtis distance algorithm was conducted to examine microbial community structure similarities between samples. LEfSe analysis was employed to identify bacterial and fungal taxa with significant differences in abundance between groups, from phylum to genus level (LDA > 2, P < 0.05).

## Statistical analysis

Statistical analysis and visualization of experimental results were conducted using GraphPad Prism software (version 8.3.0). Differences between two independent samples were assessed using a two-tailed Student’s t-test. Analysis of variance among multiple groups was performed using one-way ANOVA. Survival curves were constructed using the Kaplan-Meier method. Differences were considered statistically significant when P < 0.05. Significance levels are denoted as follows: *P < 0.05, **P < 0.01, ***P < 0.001, and ****P < 0.0001.

## Data availability

Data from bulk RNA-seq, scRNA-seq, and spatial transcriptomics (SeekSpace platform, BMKMANU S1000 platform) can be accessed from the GEO database under the accession numbers GSE270426, GSE271387, GSE270427, and GSE271222, respectively.

## Authors’ contributions

ZGC, and HZY designed the study. HZY, ZLZ, and SZJ conducted the bioinformatic analyses. HZY, GRZ, GD, YXM, JYD, and JJL carried out the experiments. HZY drafted the initial version of the manuscript. HZY, GRZ, and GD analyzed the data. ZGC, GRZ, and GD supervised the study and critically revised the manuscript. All authors read and approved the final version for publication.

## SUPPLEMENTARY FIGURE LEGENDS

Figure S1. Abscesses developed by *Ncf2*^-/-^ mice in clean-grade condions. Related to Figure 1.

**(A)** Images of lung organ of SPF WT, SPF *Ncf2*^-/-^, CL WT, and CL *Ncf2*^-/-^ mice. The arrow points to visibly apparent granulomas.

Figure S2. The alteration of ROS levels in *Ncf2* deficiency mice. Related to Figure 1.

**(A)** Representative histograms and quantitative charts of mean fluorescence intensity

for ROS levels in myeloid cells (CD45^+^CD11b^+^) of CL WT and CL *Ncf2*^-/-^ mice.

Figure S3. The alteration of immune cells in BM, SP, and PB under different conditions. Related to Figure 1.

Representative scatter plots of myeloid cells (CD11b^+^Gr1^+^, CD11b+Gr1^-^) **(A)** and lymphoid cells (CD3^+^, CD19^+^) **(B)** from BM, SP, and PB. CD11b^+^Gr1^+^ for granulocytic populations; CD11b^+^Gr1^-^ for monocytic populations, CD3^+^ for T lymphocytes, CD19^+^for B lymphocytes.

Figure S4. 16SRNA sequencing revealed changes in the pulmonary bacterial communities across four conditions. Related to Figure 1.

**(A)** Alpha diversity of bacterial communities in lung tissues of SPF WT, SPF *Ncf2*^-/-^, CL WT, and CL *Ncf2*^-/-^ mice was evaluated by chao index and shannon index.

**(B)** Bacterial community abundance at the genus level in lung tissue samples from SPF WT, SPF *Ncf2*^-/-^, CL WT, and CL *Ncf2*^-/-^ mice.

**(C)** The abundance differences of the Staphylococcus across the four groups. * p < 0.05,

** p < 0.01.

Figure S5. ITS sequencing revealed the changes in the pulmonary fungal communities across four conditions. Related to Figure 1.

**(A)** Alpha diversity of fungal communities in lung tissues of SPF WT, SPF *Ncf2*^-/-^, CL WT, and CL *Ncf2*^-/-^ mice was evaluated by chao index and shannon index.

**(B)** Fungal community abundance at the genus level in lung tissue samples from SPF WT, SPF *Ncf2*^-/-^, CL WT, and CL *Ncf2*^-/-^ mice. * p < 0.05, ** p < 0.01,*** p < 0.001.

Figure S6. Expression of marker transcripts supporting cluster grouping and annotation in lung single-cell datasets. Related to Figure 2.

**(A)** Dot plot of differentially expressed genes that identify individual cell populations as labeled in Fig. 2a.

Figure S7. Gate setting used to explore pulmonary neutrophils and macrophages. Related to Figure 2.

**(A)** Representative flow cytometry gating to identify NOS2^high^ neutrophils and indicated MDMs (NOS2^high^, Arg1^high^, NOS2^high^Arg1^high^) in the lung tissues.

Figure S8. Annotation of lung-resident macrophages. Related to Figure 4.

**(A)** UMAP shows representative genes of mesenchymal macrophages and alveolar macrophages, with color codes representing standardized gene expression. IM: interstitial macrophage, AM: alveolar macrophage.

Figure S9. Genetic deficiency in *Il1r1* alleviates hyperinflammation in CGD mice.

**(A)** Expression of *Il1a* and *Il1b* in neutrophils and macrophages in the scRNA-seq datasets.

**(B)** H&E staining of lung tissues from CL WT, CL *Il1r1^-/-^*, CL *Ncf2^-/-^*, and CL

*Ncf2^-/-^Il1r1^-/-^* mice. Scale bar: 100μm.

**(A) (C)** Representative flow cytometry plots of neutrophils and macrophages from lung tissues of CL WT, CL *Il1r1^-/-^*, CL *Ncf2^-/-^*, and CL *Ncf2^-/-^Il1r1^-/-^* mice. Bar plots displaying the proportions of CD11b^+^Ly6G^+^ neutrophils, CD11b^high^F4/80^+^ MDMs, and CD11b^low^F4/80^+^ resident macrophages. Bar plots displaying the proportions of CD11b^+^Ly6G^+^ neutrophils, CD11b^high^F4/80^+^ MDMs, and CD11b^low^F4/80^+^ resident macrophages.

**(B) (D)** Proportions of myeloid and lymphoid cells in the peripheral blood (PB), spleen (SP), and bone marrow (BM) of CL WT, CL *Il1r1^-/-^*, CL *Ncf2^-/-^*, and CL *Ncf2^-/-^Il1r1^-/-^* mice, as analyzed by flow cytometry.

Figure S10. The alterations of monocytic populations. Related to Figure 6.

**(A)** Proportions of monocytic populations in the peripheral blood (PB), spleen (SP), and bone marrow (BM) of WT(DMSO), WT(4IPP), *Ncf2*^-/-^(DMSO), and *Ncf2*^-/-^(4IPP) mice, as analyzed by flow cytometry.

Figure S11. Transcription changes in BM cells of CL *Ncf2^-/-^* mice. Related to Figure 7.

**(A)** UMAPs shows lineage specific genes of hematopoietic and immune cells, with color codes representing standardized gene expression.

**(B)** The volcano plot shows transcripts differentially expressed in BM hematopoietic and immune cells from scRNA-seq data, with red dots representing significant differences.

**(C)** The Bar plot displays the expression levels of proinflammatory transcripst (*Il1b, Il6, Tnf*) in Lin^-^ cells from Bulk RNA-seq data.

Figure S12. scRNA-seq analysis elucidated the colon tissue of CGD mice.

**(A)** UMAP plots reveals the distinct colon celltypes from CL WT (n=7,169), CL *Ncf2*^-/-^ (n=7,741) mice. Three types of immune cells (T cells, B cells, macrophages), four types of stromal cells (fibroblasts, myofibroblasts, endothelial cells, mesothelial cells), five types of intestinal epithelial cells (enterocytes, TA cells, goblet cells, tuft cells, enteroendocrine cells), and a group of neural cells (glia cells) were annotated and color-coded. ILC: Transit amplifying cells.

**(B)** Stacked bar plots showing the proportion of each cell type presented in **(A)**.

Figure S13. *Morrbid* deficiency promotes apoptosis of myeloid cells and their precursors in CGD mice

(A) Bubble chart illustrating the enrichment of apoptosis inhibition pathways in up-regulated DEGs within Lin^-^ cells(Bulk RNA-seq).

(B) Bubble chart depicting the enrichment of apoptosis inhibition pathways in up-regulated DEGs within neutrophil cluster (scRNA-seq).

(C) Proportions of hematopoietic stem and progenitor cell compartments in the bone marrow of CL WT, CL *Morrbid*^-/-^, CL *Ncf2*^-/-^, and CL *Ncf2*^-/-^*Morrbid*^-/-^ mice, analyzed via flow cytometry.

(D) Proportions of monocytic populations in the peripheral blood (PB), spleen (SP), and bone marrow (BM) of CL WT, CL *Morrbid*^-/-^, CL *Ncf2*^-/-^, and CL *Ncf2*^-/-^*Morrbid*^-/-^ mice, analyzed via flow cytometry.

Figure S14. The graphical abstract reveals the mechanisms of CGD granuloma formation and the discovery of three perturbations.

## REFERENCES

1 Dinauer, M. C. Inflammatory consequences of inherited disorders affecting neutrophil function. Blood 133, 2130–2139, doi:10.1182/blood-2018-11-844563 (2019).

2 van den Berg, J. M., et al. Chronic granulomatous disease: the European experience. PloS one 4, e5234, doi:10.1371/journal.pone.0005234 (2009).

3 Marciano, B. E. et al. Common severe infections in chronic granulomatous disease. Clinical infectious diseases : an official publication of the Infectious Diseases Society of America 60, 1176–1183, doi:10.1093/cid/ciu1154 (2015).

4 Wolach, B. et al. Genotype-Phenotype Correlations in Chronic Granulomatous Disease: Insights From a Large National Cohort. Blood, doi:10.1182/blood.2023022590 (2024).

5 Magnani, A. et al. Inflammatory manifestations in a single-center cohort of patients with chronic granulomatous disease. The Journal of allergy and clinical immunology 134, 655–662.e658, doi:10.1016/j.jaci.2014.04.014 (2014).

6 Weisser, M. et al. Hyperinflammation in patients with chronic granulomatous disease leads to impairment of hematopoietic stem cell functions. The Journal of allergy and clinical immunology 138, 219–228.e219, doi:10.1016/j.jaci.2015.11.028 (2016).

7 Sobrino, S. et al. Severe hematopoietic stem cell inflammation compromises chronic granulomatous disease gene therapy. *Cell reports*. Medicine 4, 100919, doi:10.1016/j.xcrm.2023.100919 (2023).

8 de Luca, A. et al. IL-1 receptor blockade restores autophagy and reduces inflammation in chronic granulomatous disease in mice and in humans. Proceedings of the National Academy of Sciences of the United States of America 111, 3526–3531, doi:10.1073/pnas.1322831111 (2014).

9 Conrad, A. et al. Infections in Patients with Chronic Granulomatous Disease Treated with Tumor Necrosis Factor Alpha Blockers for Inflammatory Complications. Journal of clinical immunology 41, 185–193, doi:10.1007/s10875-020-00901-8 (2021).

10 Sumaiya, K., Langford, D., Natarajaseenivasan, K. & Shanmughapriya, S. Macrophage migration inhibitory factor (MIF): A multifaceted cytokine regulated by genetic and physiological strategies. Pharmacology & therapeutics 233, 108024, doi:10.1016/j.pharmthera.2021.108024 (2022).

11 Wei, Y., Kim, J., Ernits, H. & Remick, D. The Septic Neutrophil-Friend or Foe. *Shock (Augusta*, Ga.) 55, 147–155, doi:10.1097/shk.0000000000001620 (2021).

12 Kotzin, J. J. et al. The long non-coding RNA Morrbid regulates Bim and short-lived myeloid cell lifespan. Nature 537, 239–243, doi:10.1038/nature19346 (2016).

13 He, H. et al. Prioritizing risk genes as novel stratification biomarkers for acute monocytic leukemia by integrative analysis. Discover oncology 13, 55, doi:10.1007/s12672-022-00516-y (2022).

14 Nunoi, H., Rotrosen, D., Gallin, J. I. & Malech, H. L. Two forms of autosomal chronic granulomatous disease lack distinct neutrophil cytosol factors. *Science (New York*, N.Y.) 242, 1298–1301, doi:10.1126/science.2848319 (1988).

15 Kim-Howard, X. et al. Allelic heterogeneity in NCF2 associated with systemic lupus erythematosus (SLE) susceptibility across four ethnic populations. Human molecular genetics 23, 1656–1668, doi:10.1093/hmg/ddt532 (2014).

16 Zhang, T. P. et al. Association of NCF2, NCF4, and CYBA Gene Polymorphisms with Rheumatoid Arthritis in a Chinese Population. Journal of immunology research 2020, 8528976, doi:10.1155/2020/8528976 (2020).

17 Seyedmousavi, S. et al. Exogenous Stimulation of Type I Interferon Protects Mice with Chronic Granulomatous Disease from Aspergillosis through Early Recruitment of Host-Protective Neutrophils into the Lung. mBio 9, doi:10.1128/mBio.00422-18 (2018).

18 Song, Z. et al. NADPH oxidase 2 limits amplification of IL-1β-G-CSF axis and an immature neutrophil subset in murine lung inflammation. Blood advances 7, 1225–1240, doi:10.1182/bloodadvances.2022007652 (2023).

19 Nunoi, H. et al. Treatment with Polyethylene Glycol-Conjugated Fungal D-Amino Acid Oxidase Reduces Lung Inflammation in a Mouse Model of Chronic Granulomatous Disease. Inflammation 45, 1668–1679, doi:10.1007/s10753-022-01650-z (2022).

20 Farinelli, G. et al. Lentiviral Vector Gene Therapy Protects XCGD Mice From Acute Staphylococcus aureus Pneumonia and Inflammatory Response. Molecular therapy : the journal of the American Society of Gene Therapy 24, 1873–1880, doi:10.1038/mt.2016.150 (2016).

21 Sauler, M. et al. Characterization of the COPD alveolar niche using single-cell RNA sequencing. Nature communications 13, 494, doi:10.1038/s41467-022-28062-9 (2022).

22 Li, K. et al. DJ-1 governs airway progenitor cell/eosinophil interactions to promote allergic inflammation. The Journal of allergy and clinical immunology 150, 1178–1193.e1113, doi:10.1016/j.jaci.2022.03.036 (2022).

23 Hurskainen, M. et al. Single cell transcriptomic analysis of murine lung development on hyperoxia-induced damage. Nature communications 12, 1565, doi:10.1038/s41467-021-21865-2 (2021).

24 Weiss, K. et al. Barrier Housing and Gender Effects on Allergic Airway Disease in a Murine House Dust Mite Model. ImmunoHorizons 5, 33–47, doi:10.4049/immunohorizons.2000096 (2021).

25 Wen, Y. et al. TNF-α-dependent lung inflammation upregulates PD-L1 in monocyte-derived macrophages to contribute to lung tumorigenesis. FASEB journal : official publication of the Federation of American Societies for Experimental Biology 36, e22595, doi:10.1096/fj.202200434RR (2022).

26 Martinez, F. O., Sica, A., Mantovani, A. & Locati, M. Macrophage activation and polarization. Frontiers in bioscience : a journal and virtual library 13, 453–461, doi:10.2741/2692 (2008).

27 Cable, D. M. et al. Robust decomposition of cell type mixtures in spatial transcriptomics. Nature biotechnology 40, 517–526, doi:10.1038/s41587-021-00830-w (2022).

28 Pagán, A. J. & Ramakrishnan, L. The Formation and Function of Granulomas. Annual review of immunology 36, 639–665, doi:10.1146/annurev-immunol-032712-100022 (2018).

29 Gronski, T. J., Jr., et al. Hydrolysis of a broad spectrum of extracellular matrix proteins by human macrophage elastase. The Journal of biological chemistry 272, 12189–12194, doi:10.1074/jbc.272.18.12189 (1997).

30 Kambouchner, M. et al. Lymphatic and blood microvasculature organisation in pulmonary sarcoid granulomas. The European respiratory journal 37, 835–840, doi:10.1183/09031936.00086410 (2011).

31 Broderick, L. & Hoffman, H. M. IL-1 and autoinflammatory disease: biology, pathogenesis and therapeutic targeting. Nature reviews. Rheumatology 18, 448–463, doi:10.1038/s41584-022-00797-1 (2022).

32 Lang, T. et al. Macrophage migration inhibitory factor is required for NLRP3 inflammasome activation. Nature communications 9, 2223, doi:10.1038/s41467-018-04581-2 (2018).

33 Schultze, J. L., Mass, E. & Schlitzer, A. Emerging Principles in Myelopoiesis at Homeostasis and during Infection and Inflammation. Immunity 50, 288–301, doi:10.1016/j.immuni.2019.01.019 (2019).

34 Chavakis, T., Mitroulis, I. & Hajishengallis, G. Hematopoietic progenitor cells as integrative hubs for adaptation to and fine-tuning of inflammation. Nature immunology 20, 802–811, doi:10.1038/s41590-019-0402-5 (2019).

35 Yamamoto, A., Taniuchi, S., Tsuji, S., Hasui, M. & Kobayashi, Y. Role of reactive oxygen species in neutrophil apoptosis following ingestion of heat-killed Staphylococcus aureus. Clinical and experimental immunology 129, 479–484, doi:10.1046/j.1365-2249.2002.01930.x (2002).

36 Kobayashi, S. D. et al. Gene expression profiling provides insight into the pathophysiology of chronic granulomatous disease. *Journal of immunology (Baltimore*, Md. : 1950) 172, 636–643, doi:10.4049/jimmunol.172.1.636 (2004).

37 Kasahara, Y. et al. Involvement of reactive oxygen intermediates in spontaneous and CD95 (Fas/APO-1)-mediated apoptosis of neutrophils. Blood 89, 1748–1753 (1997).

38 Gardai, S. et al. Activation of SHIP by NADPH oxidase-stimulated Lyn leads to enhanced apoptosis in neutrophils. The Journal of biological chemistry 277, 5236–5246, doi:10.1074/jbc.M110005200 (2002).

39 Cai, Z. et al. Hyperglycemia cooperates with Tet2 heterozygosity to induce leukemia driven by proinflammatory cytokine-induced lncRNA Morrbid. The Journal of clinical investigation 131, doi:10.1172/jci140707 (2021).

40 Cai, Z. et al. Targeting Bim via a lncRNA Morrbid Regulates the Survival of Preleukemic and Leukemic Cells. Cell reports 31, 107816, doi:10.1016/j.celrep.2020.107816 (2020).

41 Cai, Z. et al. Inhibition of Inflammatory Signaling in Tet2 Mutant Preleukemic Cells Mitigates Stress-Induced Abnormalities and Clonal Hematopoiesis. Cell stem cell 23, 833–849.e835, doi:10.1016/j.stem.2018.10.013 (2018).

42 Zhang, W. X. & Kubes, P. NOX2: is the best defense a good offense? Blood 139, 2851–2853, doi:10.1182/blood.2022016192 (2022).

43 Bhattacharya, S. et al. Macrophage NOX2 NADPH oxidase maintains alveolar homeostasis in mice. Blood 139, 2855–2870, doi:10.1182/blood.2021015365 (2022).

44 Oliveira, A. F. B. et al. Microbiological profile in chronic granulomatous disease patients in a single Brazilian primary immunodeficiency center. Allergologia et immunopathologia 49, 217–224, doi:10.15586/aei.v49i2.82 (2021).

45 El-Mokhtar, M. A., Salama, E. H., Fahmy, E. M. & Mohamed, M. E. "Clinical Aspects of Chronic Granulomatous Disease in Upper Egypt". Immunological investigations 50, 139–151, doi:10.1080/08820139.2020.1713144 (2021).

46 Nauseef, W. M. & Borregaard, N. Neutrophils at work. Nature immunology 15, 602–611, doi:10.1038/ni.2921 (2014).

47 Eliçabe, R. J., Arias, J. L., Rabinovich, G. A. & Di Genaro, M. S. TNFRp55 modulates IL-6 and nitric oxide responses following Yersinia lipopolysaccharide stimulation in peritoneal macrophages. Immunobiology 216, 1322–1330, doi:10.1016/j.imbio.2011.05.009 (2011).

48 Tsuji, S., Taniuchi, S., Hasui, M., Yamamoto, A. & Kobayashi, Y. Increased nitric oxide production by neutrophils from patients with chronic granulomatous disease on trimethoprim-sulfamethoxazole. Nitric oxide : biology and chemistry 7, 283–288, doi:10.1016/s1089-8603(02)00110-6 (2002).

49 Tsuji, S., Iharada, A., Taniuchi, S., Hasui, M. & Kaneko, K. Increased production of nitric oxide by phagocytic stimulated neutrophils in patients with chronic granulomatous disease. Journal of pediatric hematology/oncology 34, 500–502, doi:10.1097/MPH.0b013e3182668388 (2012).

50 Ahlin, A., Lärfars, G., Elinder, G., Palmblad, J. & Gyllenhammar, H. Gamma interferon treatment of patients with chronic granulomatous disease is associated with augmented production of nitric oxide by polymorphonuclear neutrophils. Clinical and diagnostic laboratory immunology 6, 420–424, doi:10.1128/cdli.6.3.420-424.1999 (1999).

51 Evren, E., Ringqvist, E. & Willinger, T. Origin and ontogeny of lung macrophages: from mice to humans. Immunology 160, 126–138, doi:10.1111/imm.13154 (2020).

52 Maus, U. A. et al. Resident alveolar macrophages are replaced by recruited monocytes in response to endotoxin-induced lung inflammation. American journal of respiratory cell and molecular biology 35, 227–235, doi:10.1165/rcmb.2005-0241OC (2006).

53 Santos, L. D. et al. TNF-mediated alveolar macrophage necroptosis drives disease pathogenesis during respiratory syncytial virus infection. The European respiratory journal 57, doi:10.1183/13993003.03764-2020 (2021).

54 Guilliams, M. & Svedberg, F. R. Does tissue imprinting restrict macrophage plasticity? Nature immunology 22, 118–127, doi:10.1038/s41590-020-00849-2 (2021).

55 Rath, M., Müller, I., Kropf, P., Closs, E. I. & Munder, M. Metabolism via Arginase or Nitric Oxide Synthase: Two Competing Arginine Pathways in Macrophages. Frontiers in immunology 5, 532, doi:10.3389/fimmu.2014.00532 (2014).

56 Krausgruber, T. et al. Single-cell and spatial transcriptomics reveal aberrant lymphoid developmental programs driving granuloma formation. Immunity 56, 289–306.e287, doi:10.1016/j.immuni.2023.01.014 (2023).

57 Chan, T. Y. et al. Increased ILC3s associated with higher levels of IL-1β aggravates inflammatory arthritis in mice lacking phagocytic NADPH oxidase. European journal of immunology 49, 2063–2073, doi:10.1002/eji.201948141 (2019).

58 Bagaitkar, J. et al. NADPH oxidase controls neutrophilic response to sterile inflammation in mice by regulating the IL-1α/G-CSF axis. Blood 126, 2724–2733, doi:10.1182/blood-2015-05-644773 (2015).

59 Wang, L. et al. Tumor-associated macrophages facilitate oral squamous cell carcinomas migration and invasion by MIF/NLRP3/IL-1β circuit: A crosstalk interrupted by melatonin. Biochimica et biophysica acta. Molecular basis of disease 1869, 166695, doi:10.1016/j.bbadis.2023.166695 (2023).

60 Shin, M. S. et al. Macrophage Migration Inhibitory Factor Regulates U1 Small Nuclear RNP Immune Complex-Mediated Activation of the NLRP3 Inflammasome. *Arthritis & rheumatology (Hoboken*, N.J.) 71, 109–120, doi:10.1002/art.40672 (2019).

