## Supplementary figures for "Single-cell and spatial transcriptomics reveal the pathogenesis of chronic granulomatous disease in a natural model"

### Figure S1

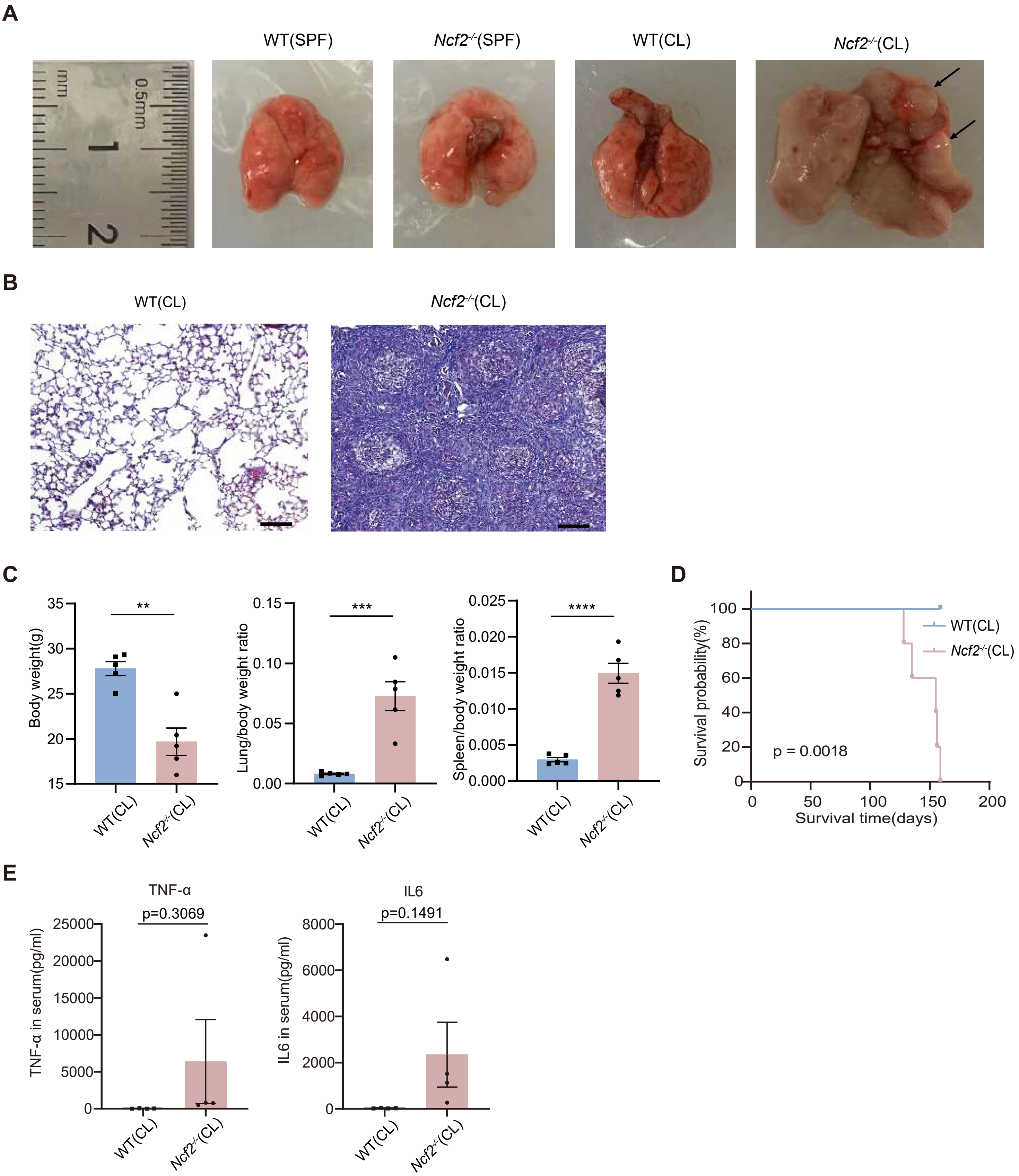
A
WT(SPF)
Ncf2-/-(SPF)
WT(CL)
Ncf2-/-(CL)
mm
0.5mm
1
2
B
WT(CL)
Ncf2-/-(CL)
C
Body weight(g)
35
30
25
20
15
**
Lung/body weight ratio
0.15
0.10
0.05
0.00
***
Spleen/body weight ratio
0.025
0.020
0.015
0.010
0.005
0.000
****
WT(CL)
Ncf2-/-(CL)
D
Survival probability(%)
100
80
60
40
20
0
WT(CL)
Ncf2-/-(CL)
p = 0.0018
0
50
100
150
200
Survival time(days)
E
TNF-α
p=0.3069
TNF-α in serum(pg/ml)
25000
20000
15000
10000
5000
0
IL6
p=0.1491
IL6 in serum(pg/ml)
8000
6000
4000
2000
0
WT(CL)
Ncf2-/-(CL)

#### Figure S2

**A**

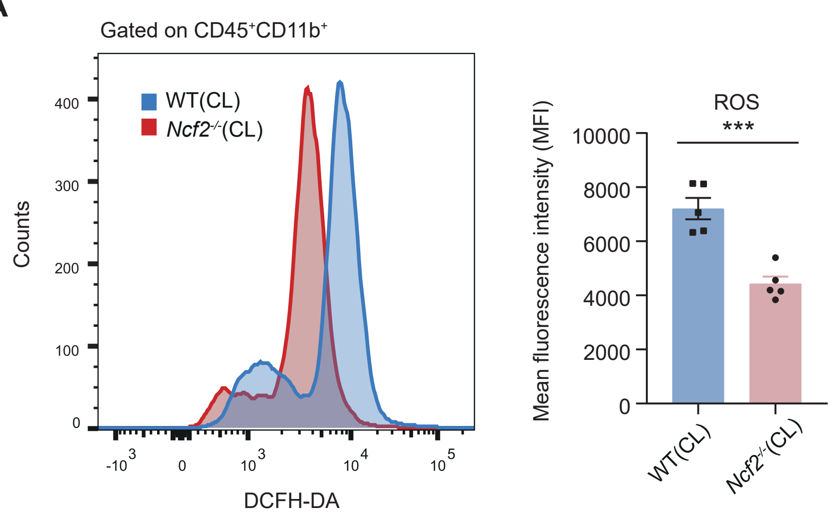

Gated on CD45+CD11b+
WT(CL)
Ncf2-/-(CL)
Counts
400
300
200
100
0
-10 3
0
10 3
10 4
10 5
DCFH-DA
ROS
***
Mean fluorescence intensity (MFI)
10000
8000
6000
4000
2000
0
WT(CL)
Ncf2-/-(CL)

**Figure S3**

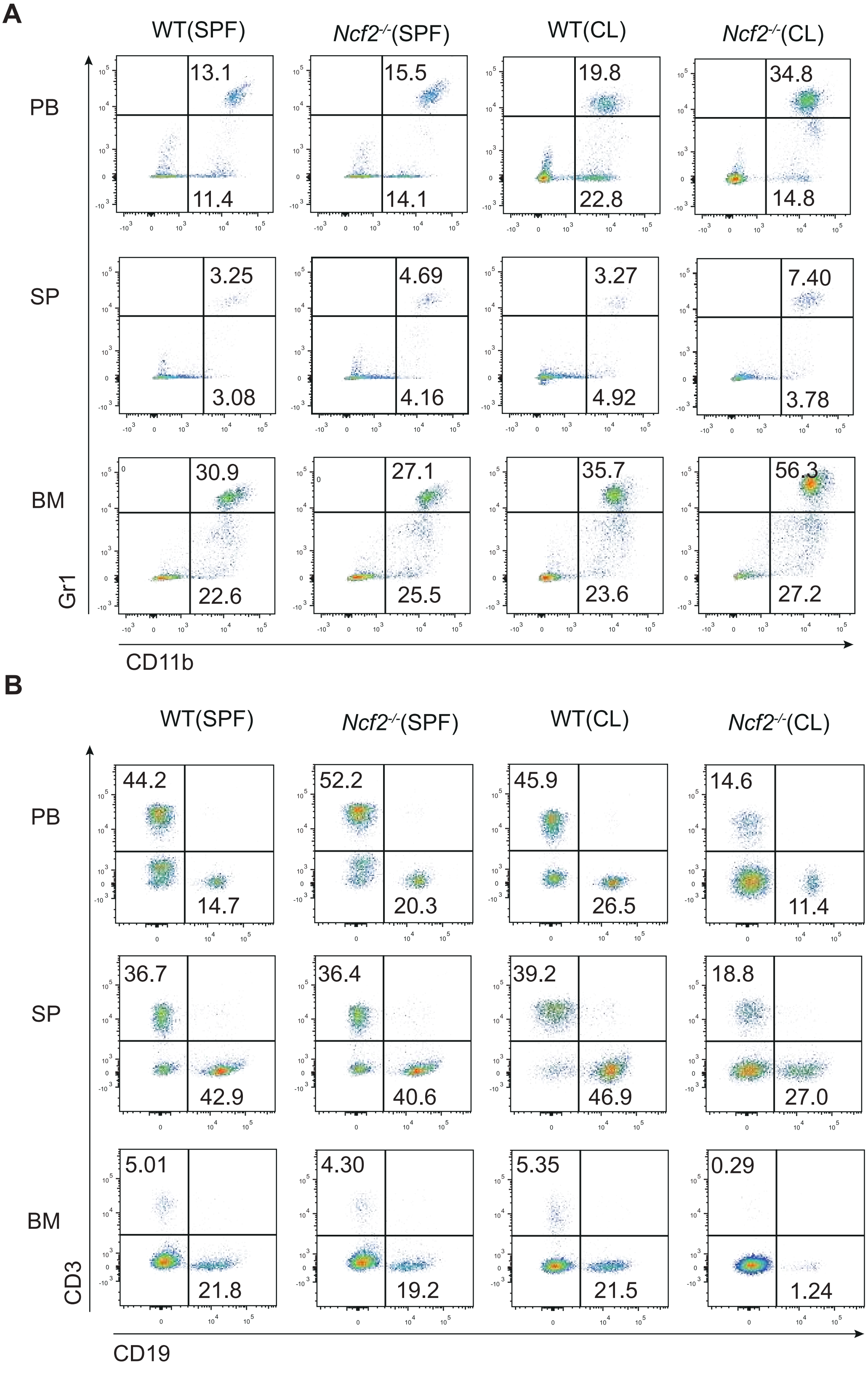
A
WT(SPF)
Ncf2-/-(SPF)
WT(CL)
Ncf2-/-(CL)
PB
13.1
11.4
15.5
14.1
19.8
22.8
34.8
14.8
SP
3.25
3.08
4.69
4.16
3.27
4.92
7.40
3.78
BM
30.9
22.6
27.1
25.5
35.7
23.6
56.3
27.2
Gr1
CD11b
B
WT(SPF)
Ncf2-/-(SPF)
WT(CL)
Ncf2-/-(CL)
PB
44.2
14.7
52.2
20.3
45.9
26.5
14.6
11.4
SP
36.7
42.9
36.4
40.6
39.2
46.9
18.8
27.0
BM
5.01
21.8
4.30
19.2
5.35
21.5
0.29
1.24
CD3
CD19

**Figure S4**

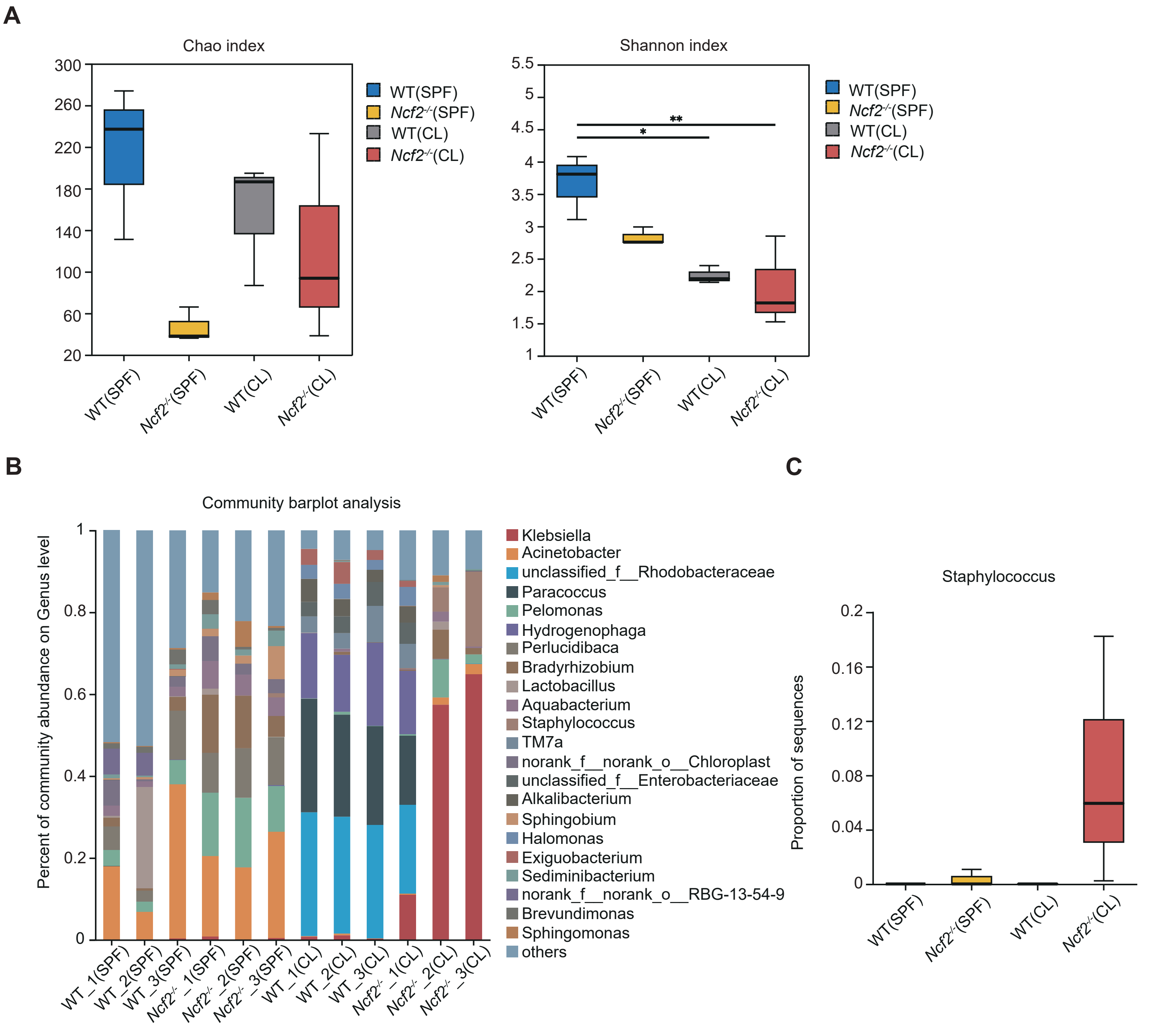
A
Chao index
WT(SPF)
Ncf2-/-(SPF)
WT(CL)
Ncf2-/-(CL)
Shannon index
**
*
B
Community barplot analysis
Percent of community abundance on Genus level
Klebsiella
Acinetobacter
unclassified_f__Rhodobacteraceae
Paracoccus
Pelomonas
Hydrogenophaga
Perlucidibaca
Bradyrhizobium
Lactobacillus
Aquabacterium
Staphylococcus
TM7a
norank_f__norank_o__Chloroplast
unclassified_f__Enterobacteriaceae
Alkalibacterium
Sphingobium
Halomonas
Exiguobacterium
Sediminibacterium
norank_f__norank_o__RBG-13-54-9
Brevundimonas
Sphingomonas
others
WT_1(SPF)
WT_2(SPF)
WT_3(SPF)
Ncf2-/- _1(SPF)
Ncf2-/- _2(SPF)
Ncf2-/- _3(SPF)
WT_1(CL)
WT_2(CL)
WT_3(CL)
Ncf2-/- _1(CL)
Ncf2-/- _2(CL)
Ncf2-/- _3(CL)
C
Staphylococcus
Proportion of sequences

#### Figure S5

**A**

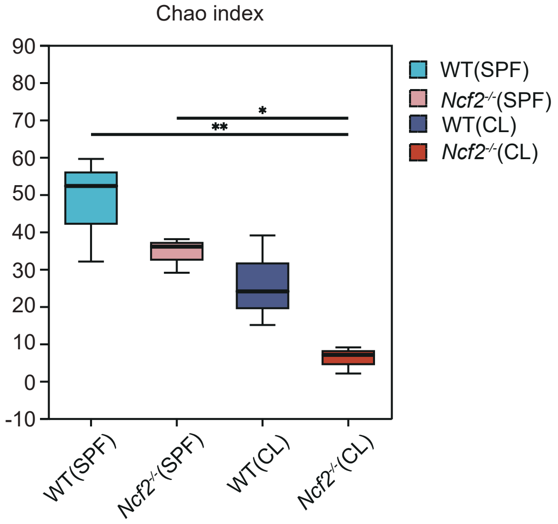
Chao index
90
80
70
60
50
40
30
20
10
0
-10
*
**
WT(SPF)
Ncf2-/-(SPF)
WT(CL)
Ncf2-/-(CL)
WT(SPF)
Ncf2-/-(SPF)
WT(CL)
Ncf2-/-(CL)

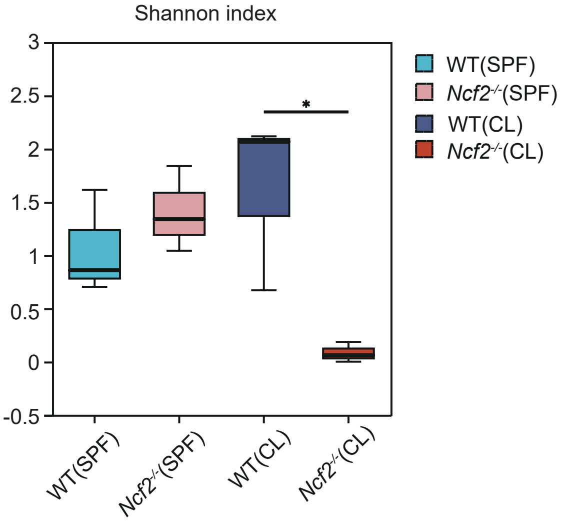
Shannon index
3
2.5
2
1.5
1
0.5
0
-0.5
*
WT(SPF)
Ncf2-/-(SPF)
WT(CL)
Ncf2-/-(CL)
WT(SPF)
Ncf2-/-(SPF)
WT(CL)
Ncf2-/-(CL)

**B**

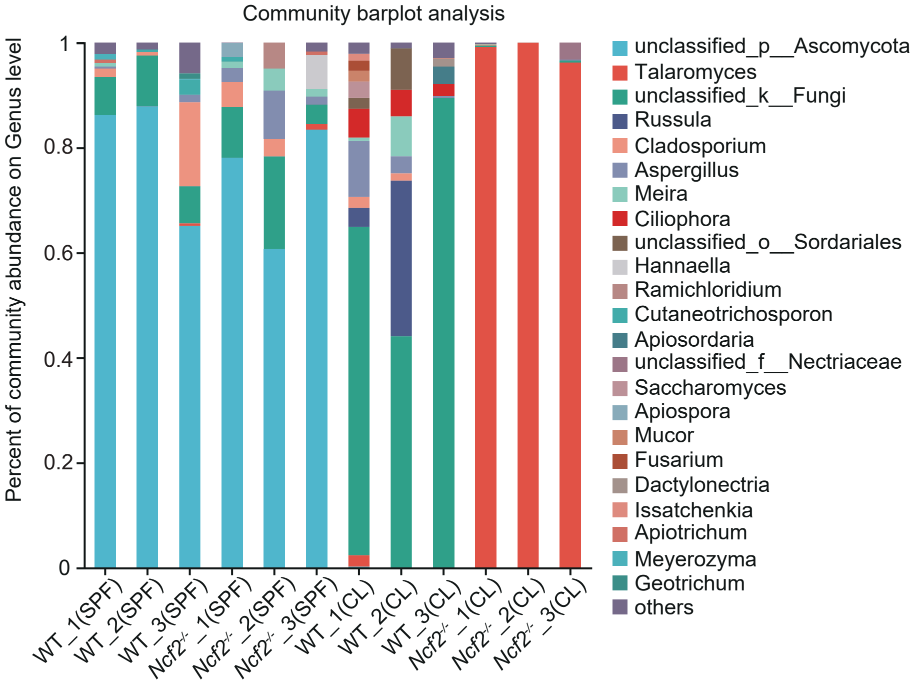
Community barplot analysis
Percent of community abundance on Genus level
1
0.8
0.6
0.4
0.2
0
WT_1(SPF)
WT_2(SPF)
WT_3(SPF)
Ncf2-/-_1(SPF)
Ncf2-/-_2(SPF)
Ncf2-/-_3(SPF)
WT_1(CL)
WT_2(CL)
WT_3(CL)
Ncf2-/-_1(CL)
Ncf2-/-_2(CL)
Ncf2-/-_3(CL)
unclassified_p__Ascomycota
Talaromyces
unclassified_k__Fungi
Russula
Cladosporium
Aspergillus
Meira
Ciliophora
unclassified_o__Sordariales
Hannaella
Ramichloridium
Cutaneotrichosporon
Apiosordaria
unclassified_f__Nectriaceae
Saccharomyces
Apiospora
Mucor
Fusarium
Dactylonectria
Issatchenkia
Apiotrichum
Meyerozyma
Geotrichum
others

#### Figure S6

**A**

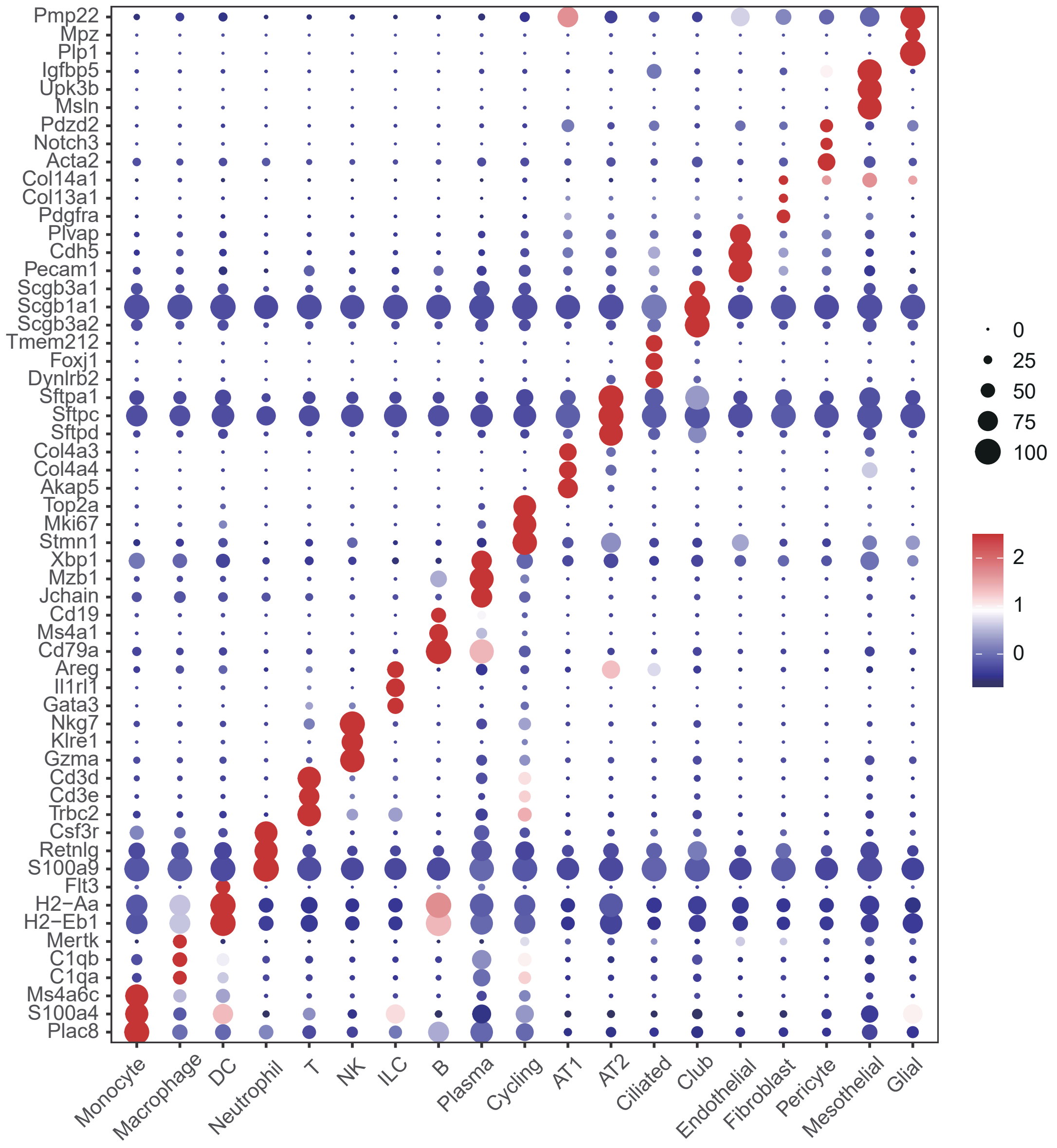

Pmp22
Mpz
Plp1
Igfbp5
Upk3b
Msln
Pdzd2
Notch3
Acta2
Col14a1
Col13a1
Pdgfra
Plvap
Cdh5
Pecam1
Scgb3a1
Scgb1a1
Scgb3a2
Tmem212
Foxj1
Dynlrb2
Sftpa1
Sftpc
Sftpd
Col4a3
Col4a4
Akap5
Top2a
Mki67
Stmn1
Xbp1
Mzb1
Jchain
Cd19
Ms4a1
Cd79a
Areg
Il1rl1
Gata3
Nkg7
Klre1
Gzma
Cd3d
Cd3e
Trbc2
Csf3r
Retnlg
S100a9
Flt3
H2−Aa
H2−Eb1
Mertk
C1qb
C1qa
Ms4a6c
S100a4
Plac8
Monocyte
Macrophage
DC
Neutrophil
T
NK
ILC
B
Plasma
Cycling
AT1
AT2
Ciliated
Club
Endothelial
Fibroblast
Pericyte
Mesothelial
Glial
0
25
50
75
100
2
1
0

### Figure S7

## A

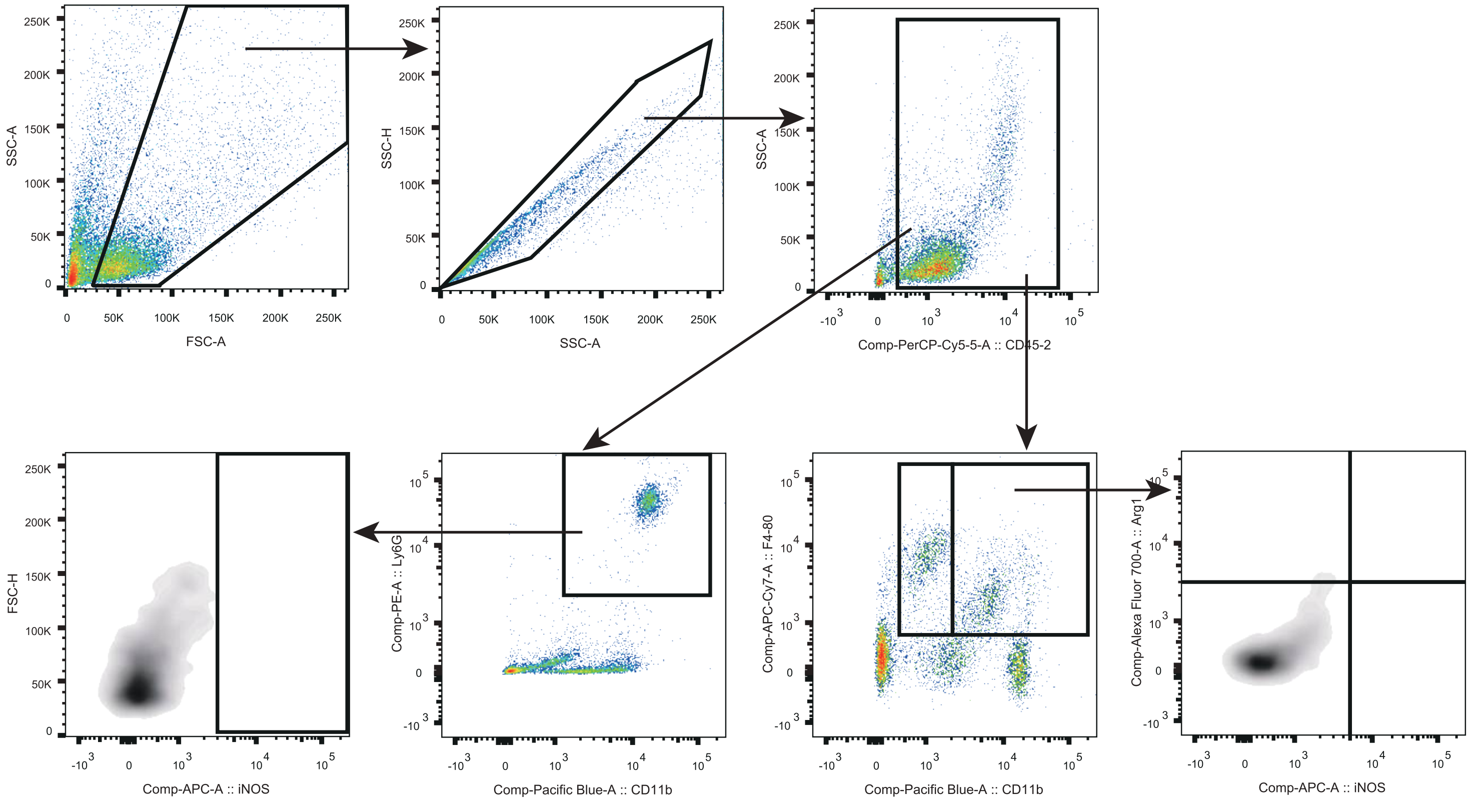

SSC-A
FSC-A
SSC-H
SSC-A
SSC-A
Comp-PerCP-Cy5-5-A :: CD45-2
FSC-H
Comp-APC-A :: iNOS
Comp-PE-A :: Ly6G
Comp-Pacific Blue-A :: CD11b
Comp-APC-Cy7-A :: F4-80
Comp-Pacific Blue-A :: CD11b
Comp-Alexa Fluor 700-A :: Arg1
Comp-APC-A :: iNOS

**Figure S8**

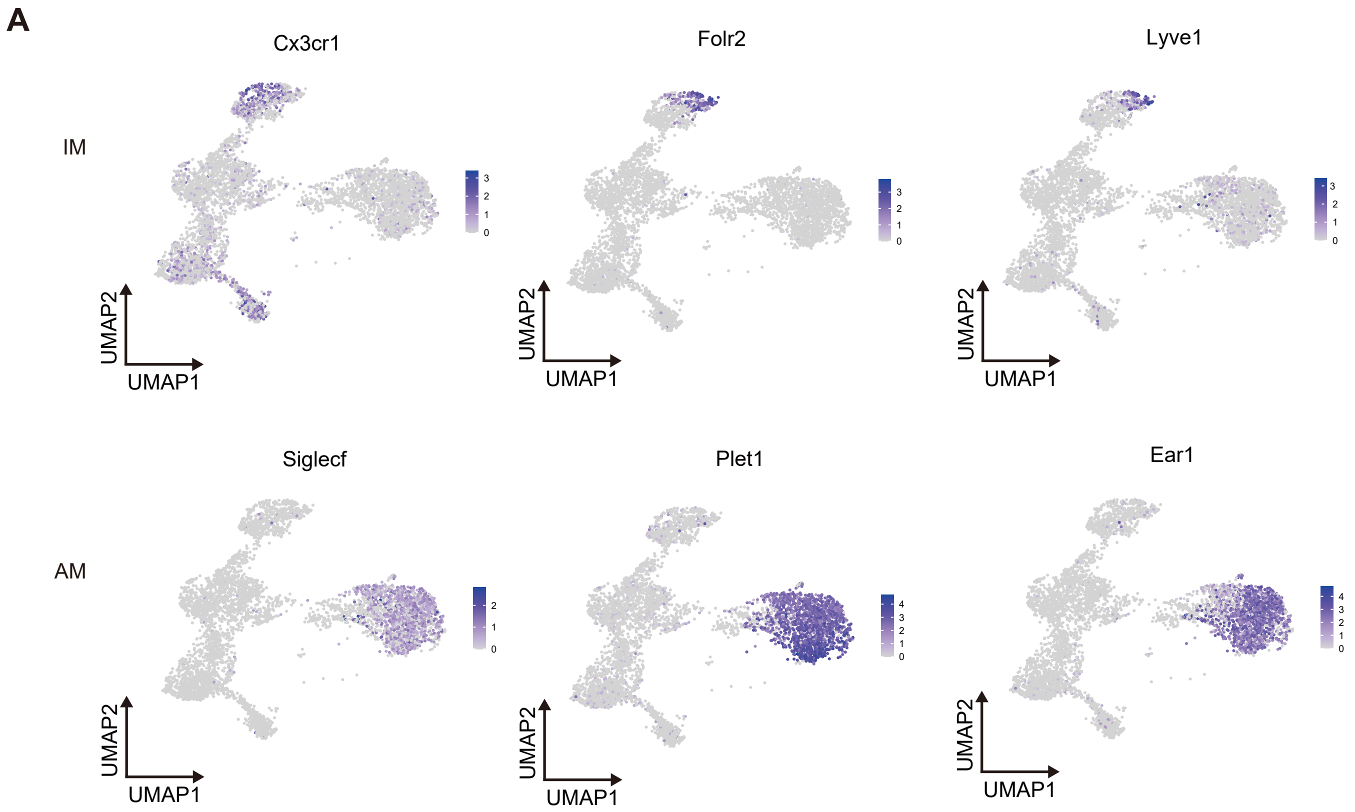

#### Figure S9

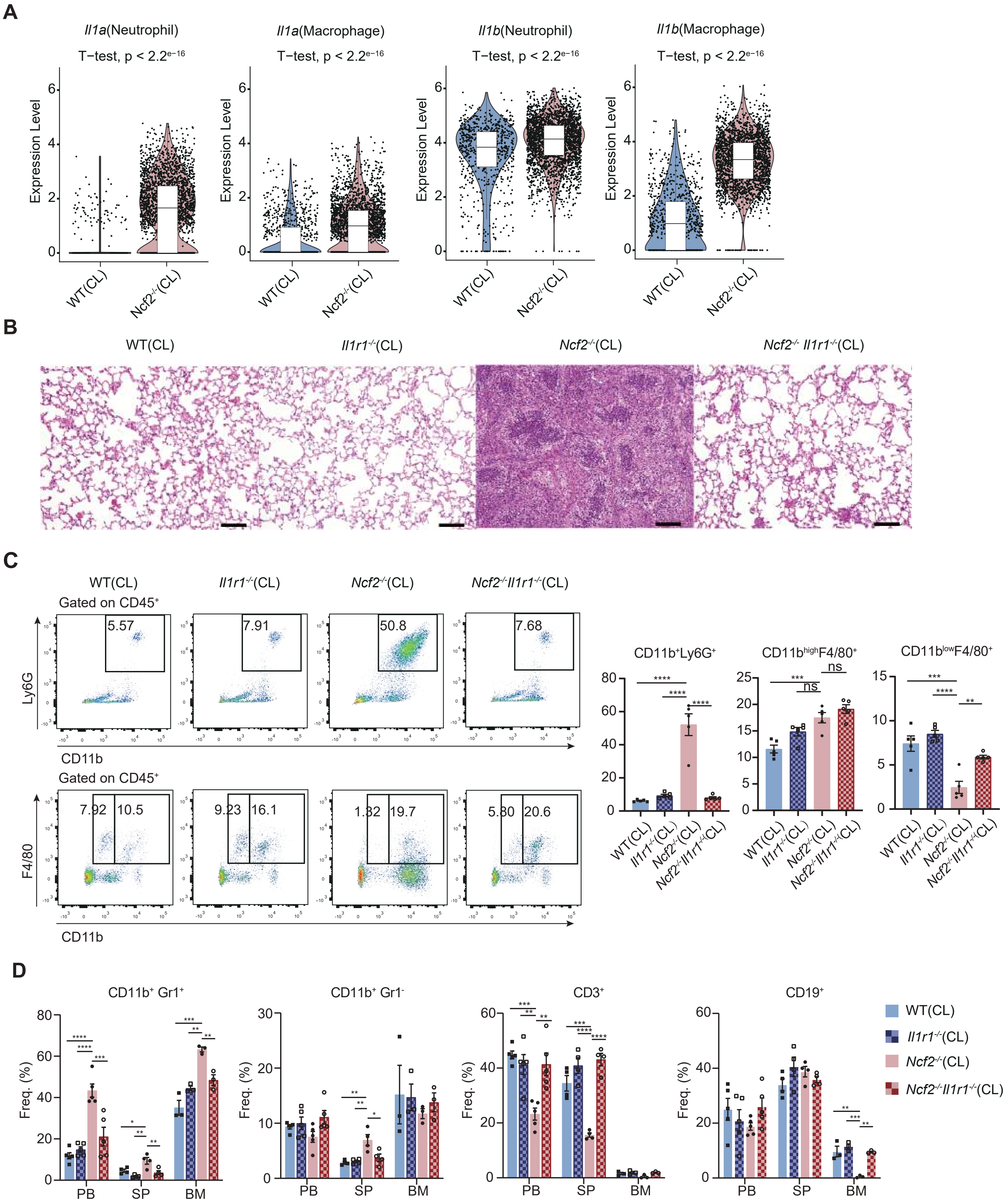

A
Il1a(Neutrophil)
Il1a(Macrophage)
Il1b(Neutrophil)
Il1b(Macrophage)
T−test, p < 2.2e−16
Expression Level
WT(CL)
Ncf2-/-(CL)
B
WT(CL)
Il1r1-/-(CL)
Ncf2-/-(CL)
Ncf2-/- Il1r1-/-(CL)
C
Gated on CD45+
Ly6G
CD11b
F4/80
5.57
7.91
50.8
7.68
7.92
10.5
9.23
16.1
1.32
19.7
5.80
20.6
CD11b+Ly6G+
CD11bhighF4/80+
CD11blowF4/80+
ns
D
CD11b+ Gr1+
CD11b+ Gr1-
CD3+
CD19+
Freq. (%)
PB
SP
BM
WT(CL)
Il1r1-/-(CL)
Ncf2-/-(CL)
Ncf2-/-Il1r1-/-(CL)

### Figure S10

## A

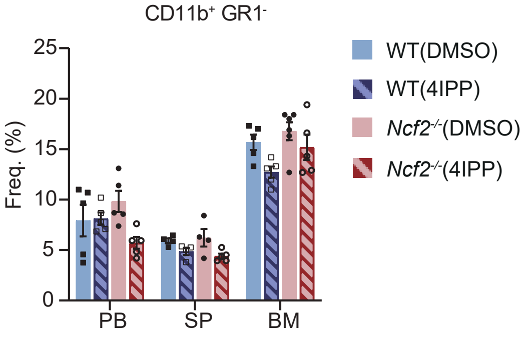

CD11b⁺ GR1⁻
Freq. (%)
25
20
15
10
5
0
PB
SP
BM
WT(DMSO)
WT(4IPP)
Ncf2-/-(DMSO)
Ncf2-/-(4IPP)

### Figure S11

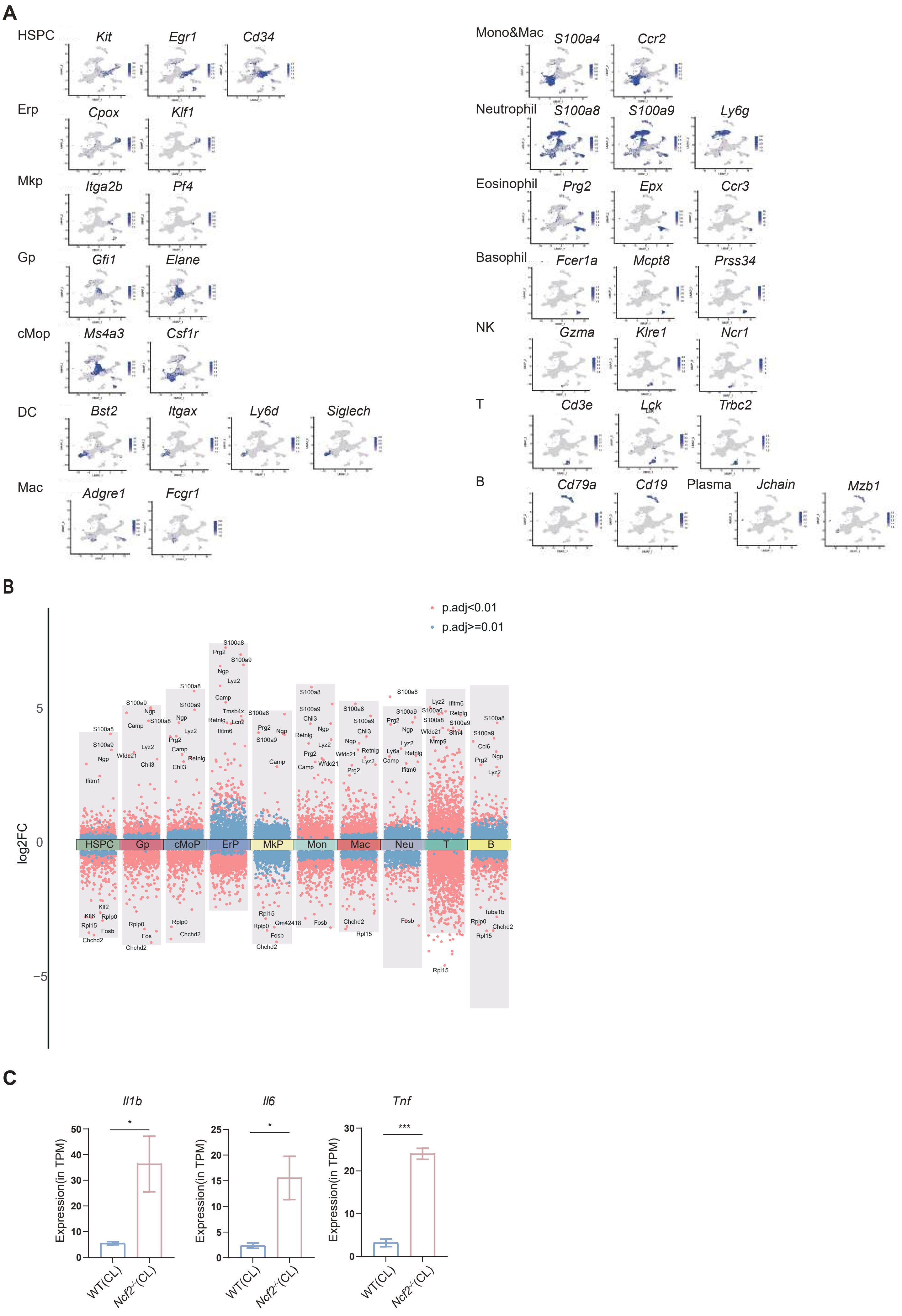
A
HSPC
Kit
Egr1
Cd34
Erp
Cpox
Klf1
Mkp
Itga2b
Pf4
Gp
Gfi1
Elane
cMop
Ms4a3
Csf1r
DC
Bst2
Itgax
Ly6d
Siglech
Mac
Adgre1
Fcgr1
Mono&Mac
S100a4
Ccr2
Neutrophil
S100a8
S100a9
Ly6g
Eosinophil
Prg2
Epx
Ccr3
Basophil
Fcer1a
Mcpt8
Prss34
NK
Gzma
Klre1
Ncr1
T
Cd3e
Lck
Trbc2
B
Cd79a
Cd19
Plasma
Jchain
Mzb1
B
p.adj<0.01
p.adj>=0.01
log2FC
5
0
−5
HSPC
Gp
cMoP
ErP
MkP
Mon
Mac
Neu
T
B
C
Il1b
Il6
Tnf
Expression(in TPM)
WT(CL)
Ncf2-/-(CL)
*
*
***

**Figure S12**

**A**

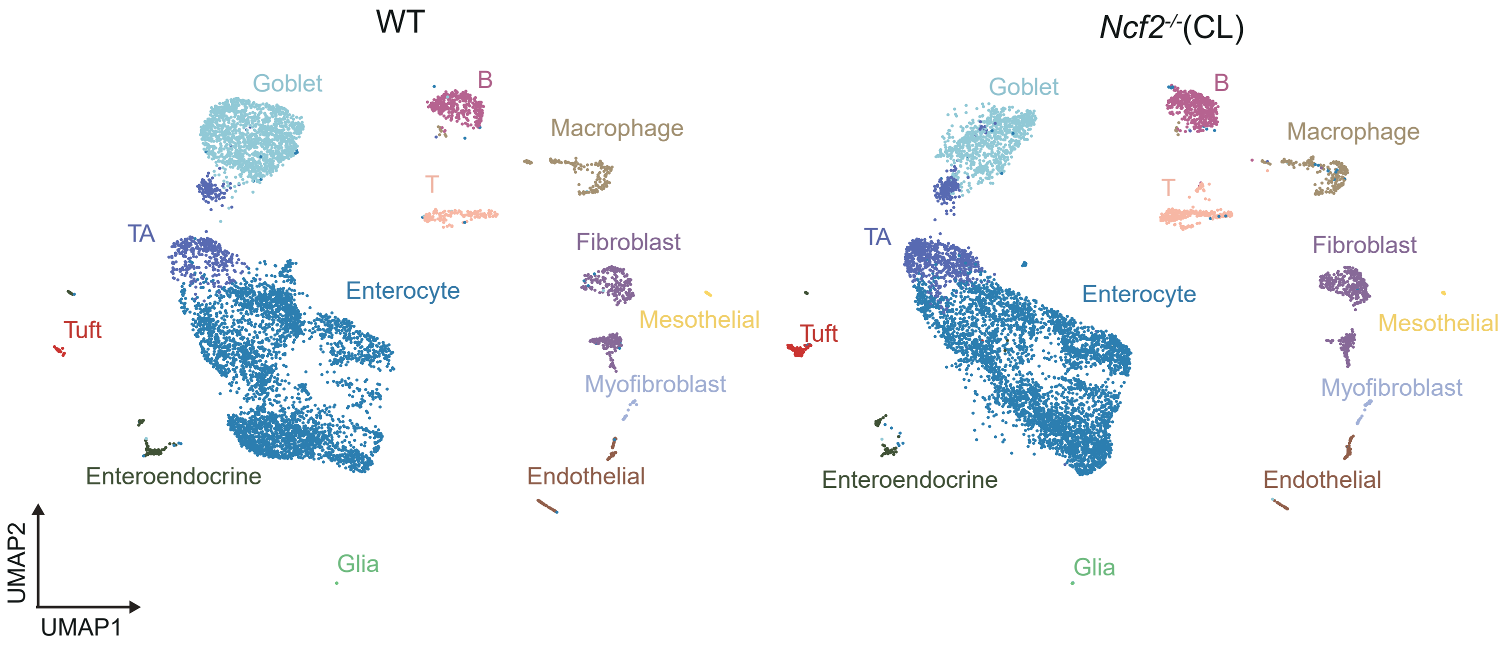


**B**

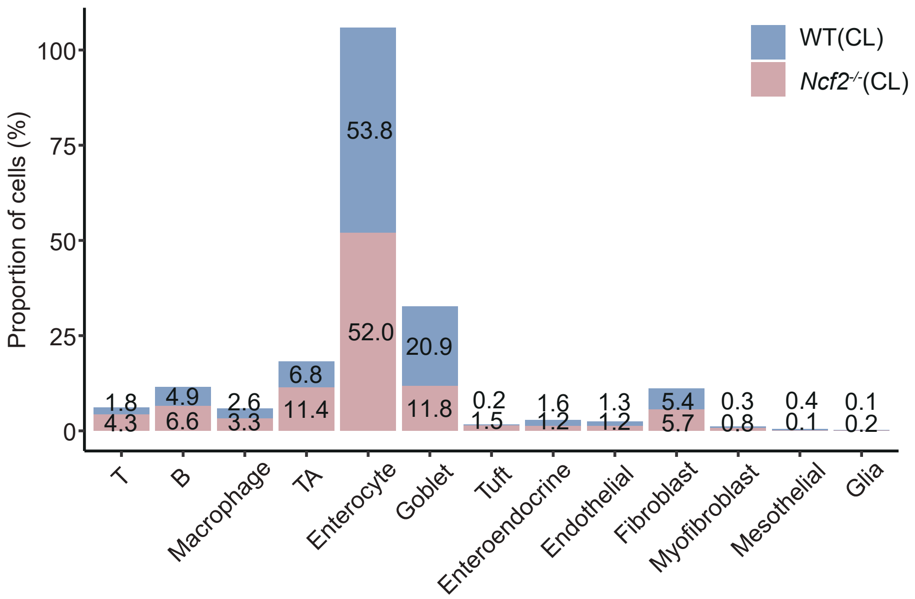

### Figure S13

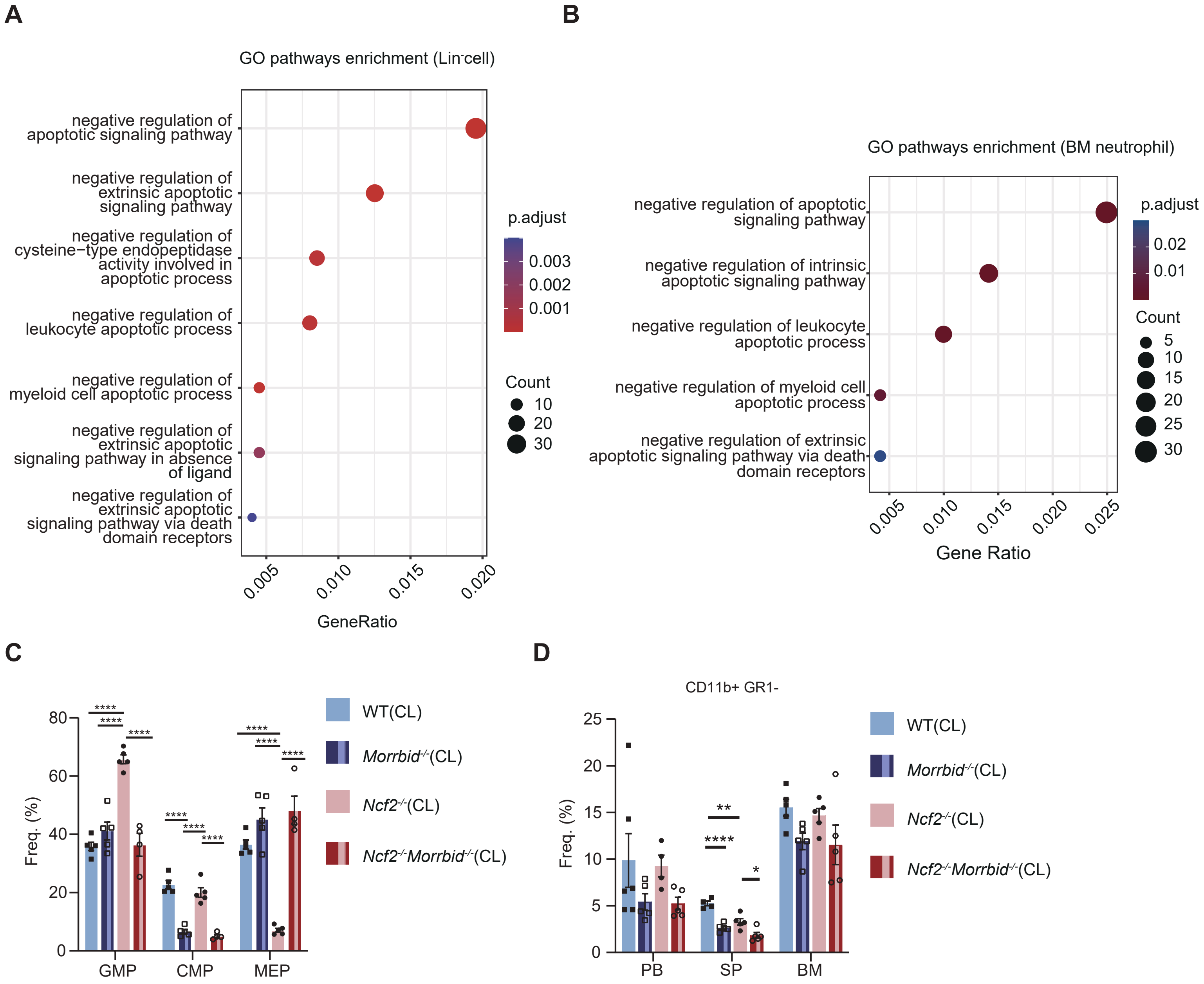
A
GO pathways enrichment (Lin-cell)
negative regulation of apoptotic signaling pathway
negative regulation of extrinsic apoptotic signaling pathway
negative regulation of cysteine−type endopeptidase activity involved in apoptotic process
negative regulation of leukocyte apoptotic process
negative regulation of myeloid cell apoptotic process
negative regulation of extrinsic apoptotic signaling pathway in absence of ligand
negative regulation of extrinsic apoptotic signaling pathway via death domain receptors
0.005
0.010
0.015
0.020
GeneRatio
p.adjust
0.003
0.002
0.001
Count
10
20
30
B
GO pathways enrichment (BM neutrophil)
negative regulation of apoptotic signaling pathway
negative regulation of intrinsic apoptotic signaling pathway
negative regulation of leukocyte apoptotic process
negative regulation of myeloid cell apoptotic process
negative regulation of extrinsic apoptotic signaling pathway via death domain receptors
0.005
0.010
0.015
0.020
0.025
Gene Ratio
p.adjust
0.02
0.01
Count
5
10
15
20
25
30
C
Freq. (%)
0
20
40
60
80
****
GMP
CMP
MEP
WT(CL)
Morrbid-/-(CL)
Ncf2-/-(CL)
Ncf2-/-Morrbid-/-(CL)
D
CD11b+ GR1-
Freq. (%)
0
5
10
15
20
25
**
****
*
PB
SP
BM
WT(CL)
Morrbid-/-(CL)
Ncf2-/-(CL)
Ncf2-/-Morrbid-/-(CL)

**Figure S14**

Granuloma formation in CL *Ncf2*$^{-/-}$ mice

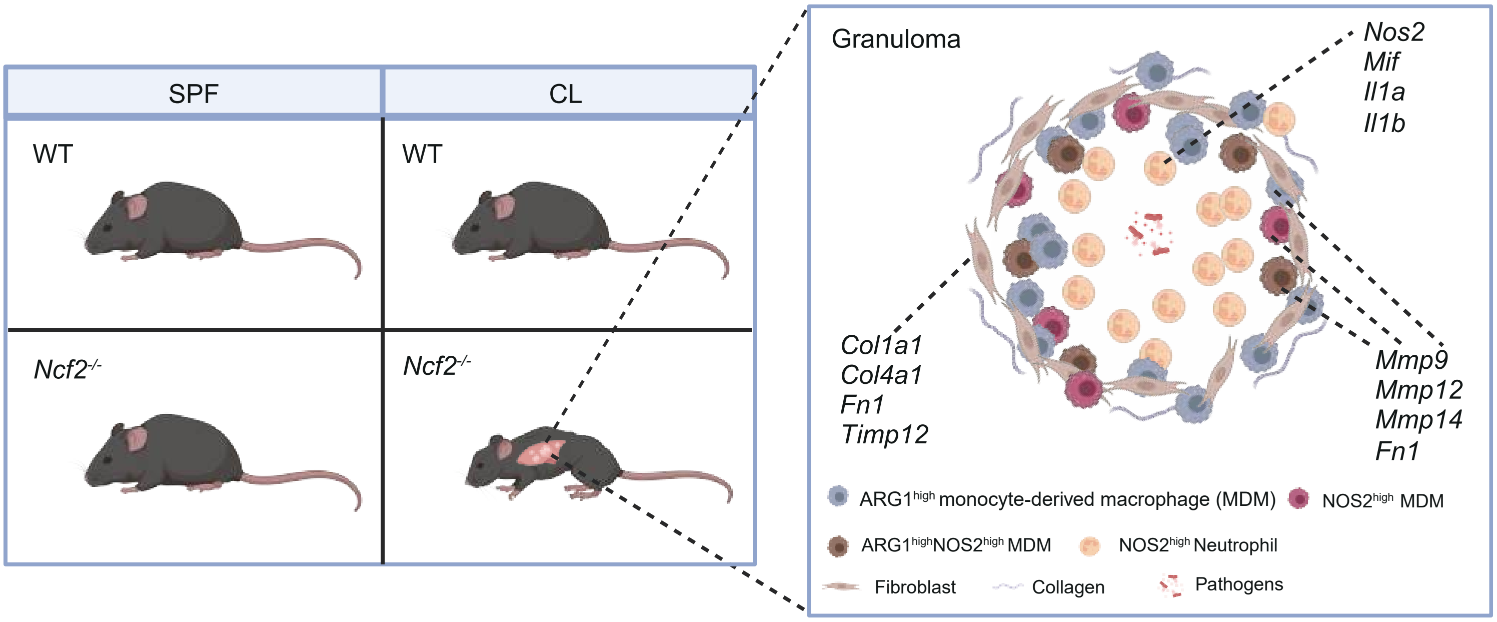


Unresolved inflammation in CGD

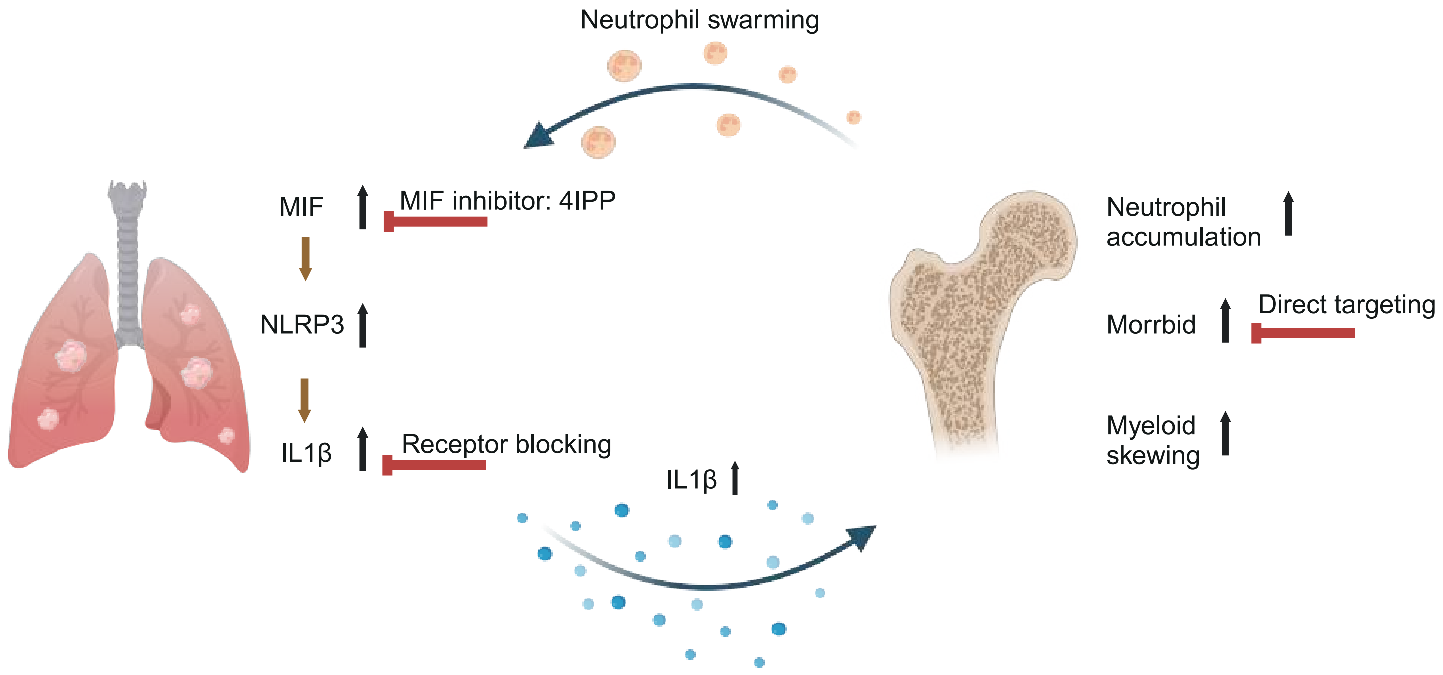
